# Representational structure of perceived food attributes in human occipitotemporal cortex

**DOI:** 10.64898/2026.03.31.715483

**Authors:** Giuseppe Marrazzo, Leonardo Pimpini, Sarah Kochs, Federico De Martino, Giancarlo Valente, Anne Roefs

## Abstract

Despite substantial progress in understanding how visual features of food are processed in the brain, it remains unclear how subjective food properties, such as perceived palatability, perceived caloric content, and perceived health value, are reflected in neural representational structure after accounting for visual similarity. Using functional MRI and representational similarity analysis (RSA), we examined visual, subjective, and categorical food representations in 25 healthy young women viewing 96 food images during an orthogonal color-discrimination task. Univariate analyses revealed reliable activation differences between high- and low-calorie foods in visual and ventral temporal cortex, with additional effects in orbitofrontal regions. Surface-based parcel-wise RSA and hypothesis-driven ROI analyses showed that visual models explained representational structure most strongly in early visual cortex and extended into occipitotemporal regions. Food-related models showed smaller but reliable effects in lateral and ventral occipitotemporal cortex: perceived calorie, perceived health, and objective calorie category were all associated with neural representational geometry in LOTC and VOTC. However, partial RSA and commonality analyses indicated that these effects were largely shared rather than separable. Perceived-calorie structure remained reliable after controlling for visual models, but not after controlling for perceived health. Palatability did not show reliable positive representational correspondence. These findings suggest that food-related information is reflected within occipitotemporal visual representational spaces during incidental viewing, but not as an isolated scalar calorie code. Instead, calorie-related effects appear to reflect a broader food-property axis on which energy density, perceived health, category structure, and learned food knowledge covary.

## Introduction

The neural representation of food is a complex and multidimensional problem, requiring the integration of visual, subjective, and nutritional information across multiple cortical systems. When viewing food, humans rapidly extract visual features such as shape, color, and texture, while simultaneously accessing higher-level information related to palatability, caloric content, and health value (Moerel, Psihoyos, & Carlson, 2024; van der Laan, de Ridder, Viergever, & Smeets, 2011; Vartanian et al., 2025). These dimensions jointly guide eating behavior and dietary choice (Rangel, 2013), yet they differ markedly in their computational demands and level of abstraction (DiCarlo, Zoccolan, & Rust, 2012; Rangel, 2013). As a result, food perception cannot be understood as a unitary process but instead likely emerges from interactions between early visual representations and higher-order systems involved in valuation, interoception, and cognitive control (Simmons, Martin, & Barsalou, 2005; Simmons et al., 2013). Understanding how these visual and subjective dimensions are represented, and how they are integrated across the cortical hierarchy, remains a central challenge in the neuroscience of food perception.

Recent research has demonstrated that visual food stimuli are processed within specialized regions of the ventral visual cortex, with food-selective responses identified in portions of the fusiform and lateral occipital cortex that are distinct from areas involved in processing faces or body parts (Bannert & Bartels, 2022; Jain et al., 2023; Khosla, Ratan Murty, & Kanwisher, 2022). However, the representational basis of this food selectivity remains debated. Some accounts emphasize visual feature statistics, including color and texture (Henderson, Tarr, & Wehbe, 2025; Pennock et al., 2023), whereas others argue that food selectivity may also reflect behaviorally relevant properties associated with eating, manipulation, reward, and learned food knowledge (Avery et al., 2025; Avery, Liu, Ingeholm, Gotts, & Martin, 2021; Simmons et al., 2005). Consistent with the latter view, a recent study showed that food and tool responses partly overlap in occipitotemporal cortex, suggesting that food representations may be shaped by graspability and action affordance (Ritchie, Andrews, Vaziri-Pashkam, & Baker, 2024). Complementarily, evidence from time-resolved multivariate EEG indicates that food representations emerge rapidly in the human brain, with early neural signals encoding food-related distinctions such as naturalness, degree of processing, and perceived caloric content shortly after stimulus onset (Moerel et al., 2024). Together, these findings suggest that food representations in visual cortex are unlikely to be organized by visual appearance alone, but instead may reflect an interaction between image-computable features and behaviorally relevant food dimensions.

Beyond perceptual processing, subjective and nutritional dimensions of food appear to play a critical role in shaping neural representations. Neuroimaging studies have consistently identified a food-related distributed cortical network that, beyond visual cortex, includes orbitofrontal cortex (OFC), the insula, and prefrontal regions (Sescousse, Caldú, Segura, & Dreher, 2013; van der Laan et al., 2011). As highlighted in recent integrative reviews (Vartanian et al., 2025), food perception cannot be reduced to visual processing alone but reflects coordinated activity across sensory, interoceptive, and valuation systems. The OFC plays a central role in encoding the subjective value of food stimuli, integrating sensory information with reward history, motivational state, and contextual factors (Kringelbach, 2005; Rolls, 2015). OFC representations of food are highly flexible and sensitive to internal state, such as hunger and satiety, as well as to learned associations and expectations (de Araujo et al., 2006; Small, Veldhuizen, Felsted, Mak, & McGlone, 2008). Importantly, OFC has been shown to encode not only overall value signals related to food, but also more specific evaluative and choice-related information, often in a distributed representational format that is not captured by univariate activation differences (Chikazoe, Lee, Kriegeskorte, & Anderson, 2014; Suzuki, Cross, & O’Doherty, 2017). This makes OFC a key candidate region for representational analyses linking subjective ratings to neural similarity structure.

The insula is critically involved in interoceptive awareness and affective evaluation of food, linking sensory representations to bodily states such as hunger, fullness, and visceral responses (Craig, 2009; Simmons et al., 2013). Food-related activity in insula has been associated with taste perception, anticipation of food intake, and integration of homeostatic signals with external food cues (Avery et al., 2021; Charbonnier et al., 2018; Small, 2010). However, the insula is a functionally heterogeneous region involved in processes well beyond food processing, including saliency detection, cognitive control, pain, and autonomic regulation (Kurth, Zilles, Fox, Laird, & Eickhoff, 2010; Menon & Uddin, 2010). Its engagement during food viewing may therefore partly reflect this broader integrative role, rather than food-specific valuation per se. While OFC is thought to encode the subjective reward value of food stimuli based on learned associations and motivational state, the insula may contribute by anchoring these evaluations in current bodily states.

Prefrontal regions, including dorsolateral prefrontal cortex (DLPFC), contribute to higher-order cognitive control, goal-directed decision-making, and regulation of food-related behavior (Hare, Camerer, & Rangel, 2009; Hutcherson, Plassmann, Gross, & Rangel, 2012). DLPFC has been implicated in self-control during food choice, modulation of value signals in OFC, and representation of long-term goals such as health considerations (Hare, Malmaud, & Rangel, 2011). From a representational perspective, DLPFC may encode more abstract, rule-based or goal-dependent aspects of food evaluation, rather than sensory or hedonic properties per se. Therefore, understanding food-related brain activity may require characterizing how representations shift from perceptual, to evaluative, to control-related formats across cortical hierarchies.

While this functional division of labor is well established, it remains unclear how these regions differ in the structure of the information they encode about food stimuli. In particular, it is not well understood how subjective evaluations relate to objective food properties at the level of neural representations. Recent work has shown that food representations are often better explained by subjective properties, such as perceived processing level or personal relevance, than by objective nutritional measures like fat or sugar content (Carrington, Liu, Candy, Martin, & Avery, 2024). This challenges traditional food categorizations based solely on macronutrient composition and highlights the importance of individual experience and cultural knowledge in food perception. These findings resonate with broader evidence that the ventral visual stream and associated networks are organized according to computational and behavioral goals rather than purely sensory inputs, with early regions encoding low-level visual features and more anterior regions representing increasingly abstract, semantic information (Grill-Spector & Weiner, 2014; Huth, Nishimoto, Vu, & Gallant, 2012; Marrazzo, De Martino, Lage-Castellanos, Vaessen, & de Gelder, 2023; Marrazzo, De Martino, Mukovskiy, Giese, & de Gelder, 2025).

While a substantial portion of neuroimaging work on food perception has relied on univariate contrasts (e.g., Killgore et al., 2003; van der Laan et al., 2011) to identify regions showing increased activation to food categories or specific attributes, these approaches have yielded mixed and often inconsistent results, with diverging findings likely stemming from heterogeneities in study design, statistical thresholding, and participant characterization (Vartanian et al., 2025). One reason for these inconsistencies may be that food-related information is encoded in the distributed structure of neural responses rather than in overall activation magnitude. Supporting this view, several multivariate fMRI studies have shown that food properties such as palatability and caloric content can be decoded above chance from multivoxel activity patterns in occipital, temporal, and frontal regions, even in the absence of univariate activation differences (Franssen, Jansen, van den Hurk, Roebroeck, & Roefs, 2020; Kochs et al., 2023; Pimpini et al., 2022), aligning with evidence that value-related information is represented in distributed neural codes (Chikazoe et al., 2014; Suzuki et al., 2017). These findings motivate the use of multivariate representational approaches to characterize how food properties are organized within neural response spaces.

Representational similarity analysis (RSA; Kriegeskorte, Mur, & Bandettini, 2008) offers a multivariate framework that enables direct testing of hypotheses about representational structure by comparing neural dissimilarity patterns to theoretically motivated model dissimilarities. RSA has been successfully applied to investigate visual object representations, semantic knowledge, and value coding across cortical systems (Kriegeskorte & Kievit, 2013; Nili et al., 2014). As the studies described above illustrate, RSA has emerged as a productive tool for dissecting food-related neural representations specifically, with recent work applying it to both EEG and fMRI data using food-relevant and visual model RDMs (Moerel et al., 2024; Ritchie et al., 2024). Using a data-driven clustering approach applied to food-responsive brain regions, a prefrontal network showed significant representational correspondence with a behavioral dimension reflecting food healthfulness and degree of processing (Avery et al., 2025). A complementary searchlight RSA in the same study further identified this correspondence in ventral occipitotemporal and parahippocampal cortex, including bilateral fusiform gyrus and lateral occipital cortex. Together, these studies establish that subjective and semantic food dimensions are recoverable from neural representational geometry.

What remains less clear, however, is how perceived food properties are expressed within specific occipitotemporal subregions, and whether closely related dimensions such as perceived calorie, perceived health, and objective calorie category explain separable or shared representational structure. This question is particularly important because recent work suggests that food selectivity in ventral visual cortex may be strongly shaped by visual features, especially color, with possible cross-modal contributions from the insula (Henderson et al., 2025). Thus, testing food-property representations requires not only comparing neural RDMs with behavioral and categorical models, but also explicitly accounting for visual similarity.

To meet this challenge, RSA can be complemented by the use of computational models of visual processing, which provide a principled way to characterize visual stimuli. Models of early visual processing (e.g. Gabor filters) offer biologically motivated approximations of orientation- and spatial-frequency–selective responses in primary visual cortex (Jones & Palmer, 1987; Kay, Naselaris, Prenger, & Gallant, 2008). Complementarily, deep convolutional neural network models optimized for object recognition, such as CORnet, have been shown to capture representational properties of mid- and high-level stages of the primate ventral visual stream, providing a computational account of hierarchical visual feature representations beyond early vision (Kubilius et al., 2019). Together, these models enable us to test whether food-property effects in occipitotemporal cortex persist beyond image-computable visual similarity.

Beyond visual models, behavioral ratings of palatability, perceived caloric content, and perceived health value can help capture subjective, graded evaluations shaped by individual experience and knowledge. Objective categorical distinctions, such as high versus low calorie content or sweet versus savory foods, provide complementary models of coarser food-category structure beyond individual subjective appraisal (Carlson, Ritchie, Kriegeskorte, Durvasula, & Ma, 2014; Clarke & Tyler, 2014; Connolly et al., 2012; Op de Beeck, Haushofer, & Kanwisher, 2008). In the present study, we combined stimulus-level fMRI modeling with RSA to investigate how different dimensions of food information are represented across the cortical hierarchy. Participants viewed a large set of food images during fMRI scanning and subsequently rated each stimulus along multiple dimensions. Using stimulus-specific beta estimates, we examined representational structure using whole-cortex Glasser parcel-wise RSA and complementary summaries within a set of a priori regions of interest spanning early visual cortex, ventral visual areas, and higher-order and control regions. Neural representational dissimilarity matrices were compared to multiple model RDMs capturing low/high-level visual similarity, subjective behavioral evaluations, and objective categorical distinctions.

The goals of this study were twofold. First, we tested whether subjective food-property judgments and objective food-category structure are reflected in neural representational geometry beyond the tested low- and high-level visual feature models. Second, we investigated how visual, subjective, and categorical food dimensions are expressed across the cortical hierarchy, with a particular focus on whether occipitotemporal food-property effects reflect separable dimensions or shared representational structure.

## Material and Methods

### Participants

Thirty-three female participants (n = 33) volunteered for this study. Only female participants were recruited to reduce variability associated with reported sex differences in neural responses to visual food cues (Chao et al., 2017; Legget et al., 2018). Participants were recruited via paper flyers, social media, and the Maastricht University student recruitment system (SONA). One participant was excluded because they experienced sickness or claustrophobia during MRI scanning and could not complete the experiment.

In addition, seven participants were excluded due to a technical error during the acquisition phase, whereby functional images were acquired using a different slice acquisition and phase-encoding direction compared to the rest of the sample. Specifically, these participants were scanned with a left–right phase-encoding direction, whereas the remaining participants were scanned with an anterior–posterior phase-encoding direction. This discrepancy resulted in substantially different susceptibility-induced distortion patterns that could not be reliably corrected or harmonized across participants. To ensure consistency of preprocessing, spatial normalization, and multivoxel pattern analyses, these participants were excluded from further analyses.

The final sample consisted of 25 participants (see Table 1 for participant characteristics). The study was pre-registered on ‘AsPredicted’ (Link) and approved by the Ethics committee of the Faculty of Psychology and Neuroscience of Maastricht University (ERCPN-252_67_04_2022). Prior participation, each participant gave informed consent and was compensated either with either 5 University credits or a 40€ voucher. At conclusion of the study, participants were compensated and received a debriefing letter.

**Table 1.**
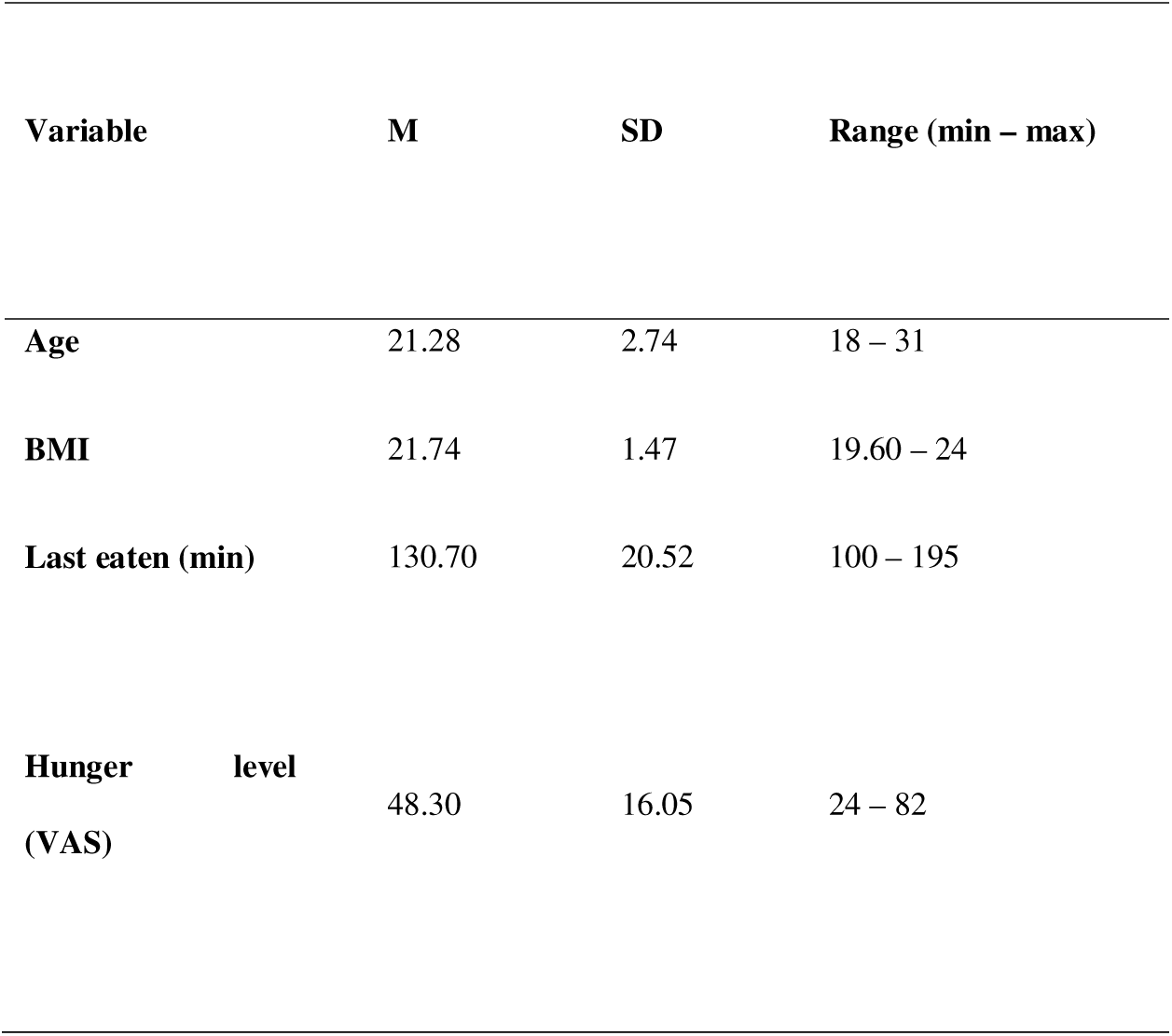
Participants’ characteristics. Reported are mean (M), standard deviation (SD), and range (minimum–maximum) for age, body mass index (BMI), time since last meal, and subjective hunger ratings. Hunger was assessed using a visual analogue scale (VAS; 0 = not hungry at all, 100 = very hungry). Time since last meal is reported in minutes.

| Variable |  | M | SD | Range (min – max) |
| --- | --- | --- | --- | --- |
| Age |  | 21.28 | 2.74 | 18 – 31 |
| BMI |  | 21.74 | 1.47 | 19.60 – 24 |
| Last eaten (min) |  | 130.70 | 20.52 | 100 – 195 |
| Hunger level (VAS) |  | 48.30 | 16.05 | 24 – 82 |

Abbreviations: min = minutes; VAS = Visual Analogue Scale (range: 0 = not hungry at all; 100 = very hungry).

### Experimental procedure

Participants completed two experimental sessions, scheduled one week apart and at a similar time of day. Approximately two weeks before the first session, they underwent a general screening for eligibility and MRI compatibility. At the start of each session, participants received instructions about scanner safety and the fMRI task and completed the hunger assessment to ensure standardized metabolic conditions (see Hunger level assessment for details).

In the second session, participants additionally completed a stimulus-rating task (∼25 minutes) after scanning (see Food rating task below), and their height and weight were measured to calculate BMI (see BMI measurement below). Each session required approximately two hours in total.

### Anthropometric and behavioral measurements

#### BMI measurement

Each participant’s height, weight, and age was measured to compute BMI (kg/m^2^; https://www.hartstichting.nl/gezond-leven/bmi). Importantly, participants were asked to take off their shoes and wear jeans/trousers and a t-shirt. The same scale and measuring tape were used throughout testing phase.

#### Hunger level assessment

To standardize metabolic state across participants, individuals were instructed to eat a small snack (e.g., a sandwich and a piece of fruit) exactly 2 hours before each session and to refrain from eating or drinking (except water) until the session ended. Upon arrival on the testing day, participants completed a hunger assessment questionnaire. This three-item paper questionnaire recorded: (1) the time elapsed since the last eating moment (in minutes), (2) a brief description of the last food consumed, and (3) subjective hunger, rated on a 100-mm visual analogue scale (0 = not hungry at all; 100 = very hungry).

#### Food rating task

Participants rated each food stimulus along four dimensions: palatability (0 = not tasty at all; 100 = very tasty), familiarity (0 = completely unfamiliar; 100 = very familiar), perceived health value (0 = very unhealthy; 100 = very healthy), and perceived caloric content (0 = very low in calories; 100 = very high in calories). Ratings were collected using a blocked design, such that all stimuli were rated on one dimension within a block. Each block was followed by a short break, after which participants proceeded to rate the stimuli on the next dimension. The order of rating blocks was randomized across participants. Within each block, food stimuli were presented individually in randomized order and rated using a 100-mm visual analogue scale (VAS). This procedure ensured that each stimulus received one rating per dimension from each participant.

#### Stimuli

In total, 96 visual food stimuli were selected from the Internet (e.g., Google, iStockphoto), and from a database issued by the University of Salzburg (Blechert, Lender, Polk, Busch, & Ohla, 2019; Blechert, Meule, Busch, & Ohla, 2014). Images were formatted using Adobe Photoshop CS5.1 to obtain a standard resolution (96 pixels/inch) and MATLAB to place each stimulus on a uniform gray background (RGB: 191, 191, 191). Stimuli subtended on average 3.7° × 2.8° (width × height) of visual angle and were presented centrally on the screen (see Figure 1A). The stimulus set was constructed to ensure an equal number of high-calorie and low-calorie food items, as well as an equal number of sweet and savory items, yielding a balanced design across these dimensions.

**Figure 1.**
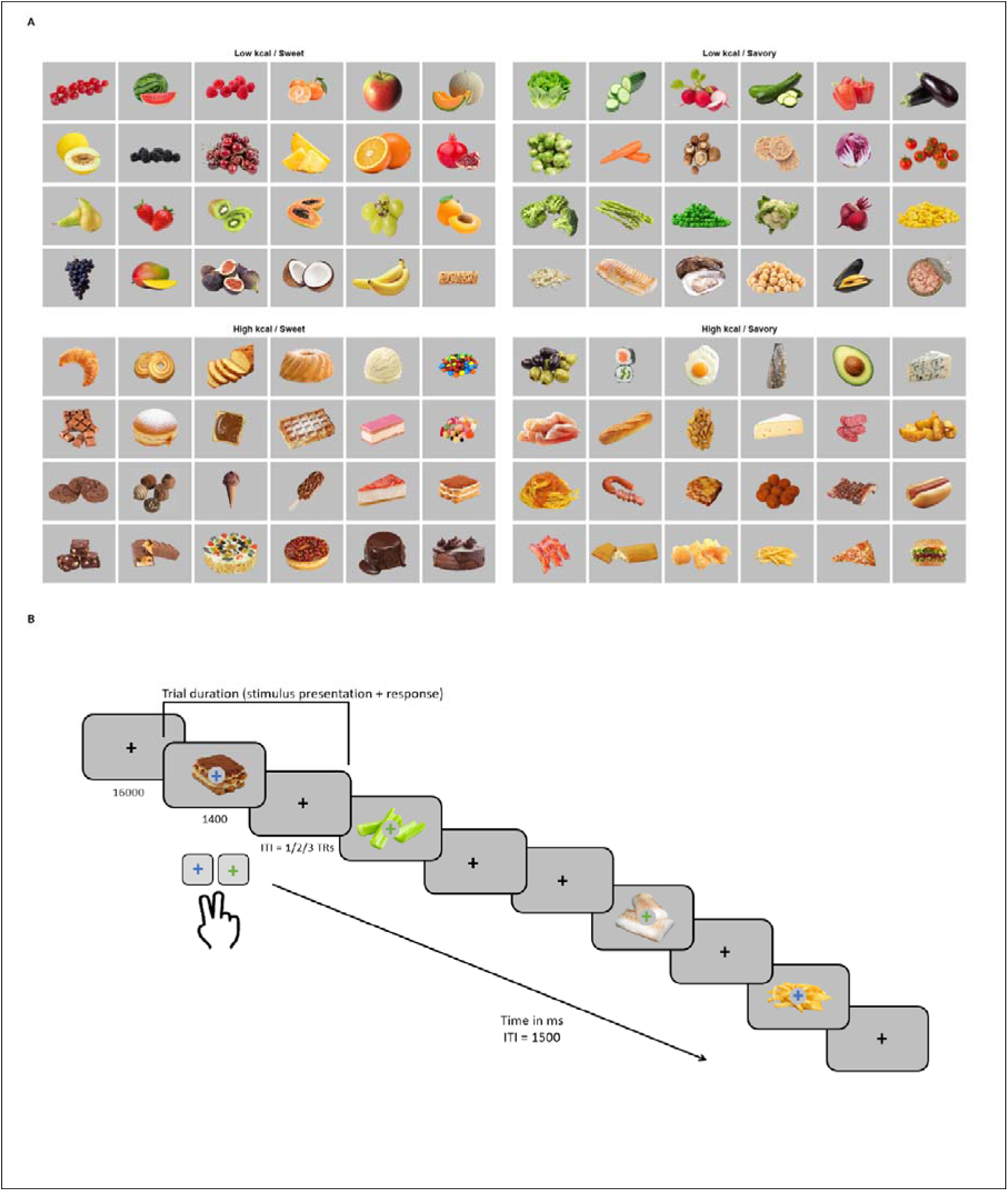
Stimuli and experimental design. (A) Stimuli were selected a priori to yield equal numbers of high- and low-calorie foods and equal numbers of sweet and savory foods. (B) Schematic illustration of the rapid event-related fMRI task. On each trial, a visual food stimulus was presented centrally for 1400 ms with a colored fixation cross (blue or green) superimposed, indicating the required response. Participants indicated the color of the fixation cross via button press. Trials were separated by a jittered inter-trial interval (ITI) of 1, 2, or 3 repetition times (TRs; TR = 1500 ms), during which a fixation cross was displayed. The order of food stimuli was randomized within each run, and null events consisting of fixation-only trials were interspersed throughout the run.

#### MRI acquisition and experimental procedure

Brain imaging was performed at Scannexus (Maastricht, The Netherlands) using a 3T Siemens MAGNETOM Prisma Fit MRI scanner (Siemens Healthineers, Erlangen, Germany) equipped with a 64-channel head–neck coil. Participants lay supine in the scanner with their head stabilized using foam padding to minimize motion. Visual stimuli were presented via a mirror mounted on the head coil.

High-resolution anatomical images were acquired using a three-dimensional T1-weighted magnetization-prepared rapid gradient-echo (MPRAGE) sequence (TR = 2250 ms, TE = 2.21 ms, inversion time = 900 ms, flip angle = 9°, field of view = 256 × 256 mm², voxel size = 1 × 1 × 1 mm³).

Functional images were acquired using a T2*-weighted multiband gradient echo–planar imaging (EPI) sequence (TR = 1500 ms, TE = 30 ms, flip angle = 71°, field of view = 208 × 208 mm², voxel size = 2 × 2 × 2 mm³). Whole-brain coverage was obtained with contiguous axial slices using multiband acceleration (factor = 3) and in-plane GRAPPA acceleration (factor = 2). Slices were acquired in an interleaved order. For each functional run, 410 volumes were collected.

Each participant completed two scanning sessions. In each session, one spin-echo EPI image with reversed phase-encoding direction was acquired and used to estimate susceptibility-induced distortions for all functional runs within that session. Two T1-weighted images acquired across sessions were combined to form a single high–signal-to-noise anatomical reference.

During functional imaging, participants performed a color-discrimination task adapted from a well-established rapid event-related fMRI paradigm (Kriegeskorte, Mur, Ruff, et al., 2008). Visual food stimuli were presented in a rapid event-related design. Each functional run (∼11 min) comprised a sequence of 96 food images, each displayed for 1.4 s, with each stimulus presented once per run. Across seven runs per session and two sessions in total, each stimulus was presented up to 14 times. Two runs were excluded across the sample for deviating volume counts, yielding on average 13.9 of the 14 runs per participant (348 runs in total).

Participants were instructed to indicate the color of a fixation cross (blue or green) superimposed on each food image by pressing a button on an MR-compatible response box. To avoid systematic response-mapping biases, fixation-cross color was pseudo-randomized across runs such that each food stimulus was paired equally often with each color (seven times with a blue cross and seven times with a green cross across the experiment). Button–color mappings were counterbalanced across participants.

Each functional run included null events to establish an implicit baseline. Four null events were presented at the beginning of each run, four at the end, and 32 were randomly interspersed among the food stimuli. Null events consisted of a centrally presented black fixation cross on a gray background. The intertrial interval was jittered and corresponded to 1, 2, or 3 repetition times (TRs).

Stimulus presentation and response collection were controlled using Presentation software (Neurobehavioral Systems, https://www.neurobs.com).

A schematic overview of the task is shown in Figure 1B.

### Preprocessing and Data analysis

MRI data were preprocessed using fMRIPrep 25.2.2 (Esteban et al., 2018; Esteban et al., 2019; RRID:SCR_016216), based on Nipype 1.10.0 (Gorgolewski et al., 2011; Gorgolewski et al., 2018; RRID:SCR_002502). The workflow included susceptibility-distortion correction, anatomical preprocessing, head-motion correction, functional-to-anatomical co-registration, cortical surface reconstruction, spatial normalization, and generation of fsLR CIFTI grayordinate outputs.

Susceptibility-induced B0 distortions were estimated from spin-echo EPI images acquired with reversed phase-encoding direction. Fourteen fieldmaps were available in the input BIDS structure, and B0-nonuniformity maps were estimated using TOPUP (Andersson, Skare, & Ashburner, 2003; FSL).

Anatomical preprocessing was performed on the two available T1-weighted images. These were corrected for intensity non-uniformity using N4BiasFieldCorrection (Tustison et al., 2010), distributed with ANTs 2.6.2 (Avants, Epstein, Grossman, & Gee, 2008, RRID:SCR_004757), combined into a single anatomical reference using mri_robust_template (FreeSurfer 7.3.2; Reuter, Rosas, & Fischl, 2010), skull-stripped, segmented into cerebrospinal fluid, white matter, and gray matter using FAST (FSL, RRID:SCR_002823; Zhang, Brady, & Smith, 2001), and used for cortical surface reconstruction with recon-all (FreeSurfer 7.3.2, RRID:SCR_001847; Dale, Fischl, & Sereno, 1999). Anatomical images were spatially normalized to MNI152NLin6Asym and MNI152NLin2009cAsym spaces using nonlinear registration with ANTs and templates accessed through TemplateFlow 25.0.4 (Ciric et al., 2022).

For each functional run, fMRIPrep generated a reference BOLD image, estimated head-motion parameters using MCFLIRT (FSL; Jenkinson, Bannister, Brady, & Smith, 2002, applied susceptibility-distortion correction, and co-registered the BOLD reference to the T1-weighted anatomical reference using boundary-based registration with bbregister (FreeSurfer; Greve & Fischl, 2009). Transformations were concatenated where possible to minimize interpolation.

For the analyses reported here, we used the fsLR CIFTI grayordinate outputs with 91k samples. Cortical data were represented on the left-right symmetric fsLR surface, and subcortical data were represented in 2-mm MNI152NLin6Asym space. Surface resampling was performed using FreeSurfer and Connectome Workbench (Glasser et al., 2013), whereas volumetric resampling was performed using nitransforms with cubic B-spline interpolation. The use of fsLR grayordinate data allowed subsequent ROI and RSA analyses to be performed in a surface-based coordinate system while retaining the full grayordinate representation generated by fMRIPrep.

fMRIPrep also generated nuisance regressors, including motion parameters, framewise displacement, DVARS, global signals, CompCor components (Behzadi, Restom, Liau, & Liu, 2007) motion outlier regressors, and temporal derivatives/quadratic expansions of selected confounds (Satterthwaite et al., 2013). Volumes exceeding 0.5 mm framewise displacement or 1.5 standardized DVARS were annotated as motion outliers. This threshold was distinct from the preregistered gross-motion exclusion criterion of 3 mm/degrees; no participant exceeded that criterion. These fMRIPrep-derived nuisance regressors were used only in the category-level GLM. They were not entered into the stimulus-level beta-estimation procedure used for RSA, which was performed with GLMsingle (Prince et al., 2022), where nuisance variance was handled within the GLMsingle framework.

### First-level GLM analysis for RSA

Stimulus-specific neural responses were estimated using GLMsingle (Prince et al., 2022), applied directly to the unsmoothed fMRIPrep-preprocessed fsLR den-91k CIFTI time series. No additional spatial smoothing was applied, because the RSA analyses were based on distributed grayordinate-level response patterns.

For each included run, a binary design matrix was constructed with 96 columns, one for each food stimulus. Stimulus events were modeled at their observed run volume, and null events were left as implicit baseline. The design matrices from all usable runs were entered jointly into GLMsingle, together with a participant-specific session indicator reflecting the retained runs from the two scanning sessions. GLMsingle was configured to estimate stimulus responses using HRF library fitting, GLMdenoise, fractional ridge regression, and percent-BOLD normalization. No separate nuisance-regression or pre-cleaning step was applied before GLMsingle; denoising was performed within the GLMsingle framework.

Trial-wise response estimates were extracted from the GLMsingle Type-D output, which combines HRF fitting, GLMdenoise, and fractional ridge regression. The resulting beta matrix contained one response estimate per available food-image presentation and one column per cortical grayordinate. A good-grayordinate mask was then defined for each participant by retaining grayordinates with finite beta values across all extracted trials. Trial-wise betas were restricted to this mask.

For each participant, trial-wise betas were averaged within stimulus identity to obtain one condition-level response pattern per image, yielding a 96 × grayordinates response matrix. These condition-level beta patterns were used for the subsequent parcel-wise and ROI-based RSA analyses.

### Category-level GLM

The univariate category-level contrast analysis was included for completeness, providing a conventional characterization of mean activation differences between food categories. To this end, a category-level general linear model (GLM) was estimated for each participant, operating on preprocessed fMRIPrep fsLR den-91k CIFTI dtseries.

Task events were defined based on stimulus category. Each food stimulus presentation was assigned to one of two caloric categories (high-calorie/low-calorie). Event onsets were modeled at the time of stimulus presentation, with a fixed stimulus duration. Task regressors were convolved with a canonical hemodynamic response function to model the expected BOLD response.

To control for non-task-related sources of variance, nuisance regressors were included comprising six rigid-body motion parameters, mean white matter and cerebrospinal fluid signals, and cosine regressors modeling low-frequency signal drifts. For each participant, retained runs were concatenated in time, with confound regressors modeled as run-specific nuisance columns and run-specific intercepts included in the design matrix. The category-level GLM was fit to the CIFTI time-series data using ordinary least squares.

For this conventional univariate category-level analysis, we smoothed fsLR-91k CIFTI time series with Connectome Workbench using a 6-mm FWHM surface kernel and no volumetric smoothing, whereas the trial-level GLM used for RSA was performed on unsmoothed CIFTI data to preserve fine-grained stimulus-level response differences.

For each participant, a contrast comparing high-calorie versus low-calorie food presentations was computed, yielding a subject-level 91k grayordinate contrast map that quantified category-related differences in mean BOLD response. A subject-level grayordinate validity mask was computed from the retained CIFTI time series by including grayordinates with finite values, non-zero signal, and non-zero temporal variance.

Group-level inference was conducted using a second-level random-effects analysis. Subject-specific first-level effect maps from the high-calorie versus low-calorie contrast were entered into a grayordinate-wise one-sample t-test across participants to assess whether the mean contrast differed significantly from zero. Multiple comparisons were controlled using Benjamini-Hochberg FDR correction across finite grayordinates for the contrast.

Importantly, regions of interest for the RSA were not defined with this analysis, instead they were defined independently, based on anatomical and functional considerations motivated by previous literature. ROI definitions were derived from the multimodal cortical parcellation proposed by Glasser and colleagues (Glasser et al., 2016), ensuring an unbiased and anatomically grounded selection of cortical regions. This approach avoided circularity and ensured that multivariate representational analyses were not constrained by univariate activation effects observed in the current dataset.

### Representational Similarity Analysis

Representational similarity analysis (RSA; Kriegeskorte, Mur, & Bandettini, 2008) was performed on condition-level beta estimates derived from the trial-level GLM to assess how neural activity patterns represent different dimensions of food stimuli. Although the pre-registered analysis plan specified whole-brain searchlight RSA, this approach did not yield results surviving correction for multiple comparisons across the whole brain. We therefore adopted a parcel-based ROI approach, which offers greater statistical sensitivity by pooling grayordinates within anatomically defined cortical parcels and reducing the number of comparisons. Analyses were conducted using bilateral parcels from the HCP-MMP1.0 atlas (Glasser et al., 2016). For visualization and hypothesis-driven ROI-level inference, parcel-wise RSA values were subsequently summarized within six a priori ROI groups: primary visual cortex (V1); lateral occipitotemporal cortex (LOTC: LO1, LO2, LO3); ventral occipitotemporal cortex (VOTC: FFC and VVC); orbitofrontal cortex (OFC: 11l, 13l, OFC, pOFC); insula (PoI2, MI, AVI, AAIC, PoI1, Ig, and PI); and dorsolateral prefrontal cortex (DLPFC: 8Av, 8Ad, 8BL, 8C, 9p, 9a, p9-46v, a9-46v, 9-46d, 46, i6-8, s6-8). These groups were chosen to span early visual, higher-order perceptual, subjective, interoceptive, and cognitive-control regions. ROI groups are shown in Figure 3A, and included parcels and valid grayordinates are listed in Supplementary Table 2.

For parcel-wise RSA, grayordinate-wise beta values were extracted for all 96 stimulus conditions within each bilateral Glasser parcel. Grayordinates with non-finite values, all-zero responses, or zero variance across conditions were excluded; all remaining valid grayordinates were retained. For each participant and parcel, neural representational dissimilarity matrices (RDMs) were computed using correlation distance, defined as 1 − Pearson correlation, between multivariate response patterns. RDMs were vectorized by extracting the upper-triangular elements and compared with model RDMs at the single-subject level using Spearman rank correlation. For ROI-level RSA, parcels belonging to the same ROI group were pooled within participant before feature cleaning/selection and neural-RDM computation, yielding one neural RDM per participant and ROI group. Subject-level RSA correlations were Fisher-z transformed for group-level inference; reported effects are shown on the correlation scale.

To contextualize RSA effect sizes, we estimated split-half neural RDM reliability for each participant and ROI. Trial-level beta estimates for each stimulus were randomly divided into two halves across repetitions, condition-average response patterns were computed separately for each half, and two neural RDMs were constructed using the same distance metric and ROI features as in the main RSA. Split-half reliability was defined as the Spearman correlation between the vectorized upper triangles of the two RDMs. This procedure was repeated 100 times with different random splits, and correlations were averaged after Fisher transformation. The mean split-half reliability was Spearman–Brown corrected to estimate full-RDM reliability and converted to an r-space noise ceiling by taking the square root of the non-negative corrected reliability. Noise ceilings are shown as mean ± SEM across participants and used only as descriptive effect-size benchmarks, not as inferential thresholds.

### Representational models and regional hypotheses

Representational models were constructed to test distinct hypotheses about the structure of neural responses to food stimuli, while dissociating low-level visual similarity from higher-level perceptual, objective, and subjective food dimensions.

Low-level visual similarity was modeled using a Gabor-based feature space approximating early visual, V1-like processing. Each stimulus image was converted to grayscale, resized to 256 × 256 pixels, standardized to zero mean and unit variance, and filtered with two-dimensional Gabor filters varying across eight orientations from 0 to π and three spatial frequencies (0.08, 0.16, and 0.32 cycles/pixel). Phase-invariant Gabor energy was computed as the magnitude of the complex-valued filter response. To preserve coarse spatial structure while allowing spatial pooling, each energy map was divided into a 4 × 4 grid, and mean energy was extracted from each cell. Pooled responses were concatenated across orientations and frequencies, z-scored, and converted into a representational dissimilarity matrix (RDM) using correlation distance. Because Gabor filters approximate orientation-and spatial-frequency–selective simple-cell responses in primary visual cortex (Jones & Palmer, 1987; Kay et al., 2008), this model served as a control for early visual similarity. We expected Gabor correspondence to be strongest in V1 and to decrease in higher-order regions.

Mid- and high-level visual representations were modeled using CORnet-S, a pretrained biologically motivated deep neural network designed to approximate stages of the primate ventral visual stream, including intermediate V4-like and higher-level IT-like representations (Kubilius et al., 2019).

CORnet models have been shown to produce representational geometries that align with neural response patterns in visual cortex, making them suitable candidate models for representational similarity analysis.

Stimulus images were processed using standard ImageNet preprocessing. Images were resized and center-cropped to 224 × 224 pixels, converted to RGB tensors, and normalized channel-wise using the ImageNet mean and standard deviation (mean = [0.485, 0.456, 0.406], SD = [0.229, 0.224, 0.225]). Images were passed through a pretrained CORnet-S network, and activations were extracted from the V4 and IT modules. For each layer, activations were stacked across stimuli to form stimulus-by-feature matrices.

Because deep networks feature spaces can be highly redundant, dimensionality reduction was performed using principal component analysis (PCA) separately for V4 and IT. PCA was applied to z-scored feature matrices (z-scoring across stimuli for each feature), retaining the number of components required to explain 99% of the variance. CORnet V4 and IT RDMs were then computed using correlation distance between stimulus feature vectors in PCA space. These models captured increasingly abstract visual similarity structure along the ventral hierarchy, including object form, texture, and compositional information. We expected CORnet correspondence to be stronger in higher-level visual cortex, including LOTC, than in early visual cortex, and used these models as high-level visual benchmarks against which subjective food-property models were evaluated.

A color-based model was constructed to capture global chromatic similarity. Images were resized to 224 × 224 pixels, RGB values were normalized to [0, 1], and images were converted to CIELAB color space. Color information was summarized using a three-dimensional histogram over the L*, a*, and b* channels. Histograms were flattened, normalized to unit sum, and converted into a color RDM using correlation distance. This model captured global hue and luminance composition while remaining agnostic to spatial layout and higher-level visual or conceptual structure. It was included to test whether ventral visual RSA effects could be attributed to systematic color statistics that covaried with food properties.

Subjective food-property models were derived from participants’ ratings of palatability, perceived caloric content, and perceived health value. Ratings were z-scored within participant to remove individual differences in scale use and response offset, then averaged across participants to obtain group-level stimulus scores. Separate group perceived-palatability, perceived-calorie, and perceived-health RDMs were constructed from pairwise absolute differences between stimuli in these group-average scores. Thus, these models captured group-level inter-stimulus dissimilarity along each subjective dimension.

Reliability of the group-level subjective models was estimated using split-half consistency across participants. Participants were randomly divided into two approximately equal groups 10,000 times. For each split, group-average rating vectors and corresponding absolute-difference RDMs were computed separately for each half, correlated using Spearman correlation, and corrected using the Spearman–Brown formula.

As a complementary control, subject-specific behavioral RDMs were also constructed from each participant’s own ratings of perceived calorie, perceived health, and palatability. For each participant and dimension, ratings were z-scored across stimuli and converted into pairwise absolute-difference RDMs. These participant-specific model RDMs were matched to the same participant’s neural RDM and analyzed separately from the group-level behavioral models, testing whether each individual’s own rating geometry predicted their own neural representational geometry.

Finally, objective categorical models were constructed for coarse food-category structure. One model distinguished high- from low-calorie foods, and another distinguished savory from sweet foods. Stimuli were assigned predefined binary labels, and dissimilarities were computed using Euclidean distance, yielding categorical RDMs in which stimuli from different categories were more dissimilar than stimuli from the same category. These models tested whether neural representations reflected categorical food structure beyond continuous subjective ratings, with expected contributions in higher-level associative regions, including OFC and DLPFC.

To visualize relationships among candidate models, we computed pairwise dissimilarities between model RDMs as 1 − Spearman correlation and summarized this model space using multidimensional scaling (MDS; Fig. 2B). This analysis was independent of neural data and served to show which visual, subjective, and categorical models captured overlapping versus distinct representational structure. Supplementary Table 3 summarizes each model, its intended interpretation, construction, and, for control analyses, the comparison tested relative to the main effects.

**Figure 2.**
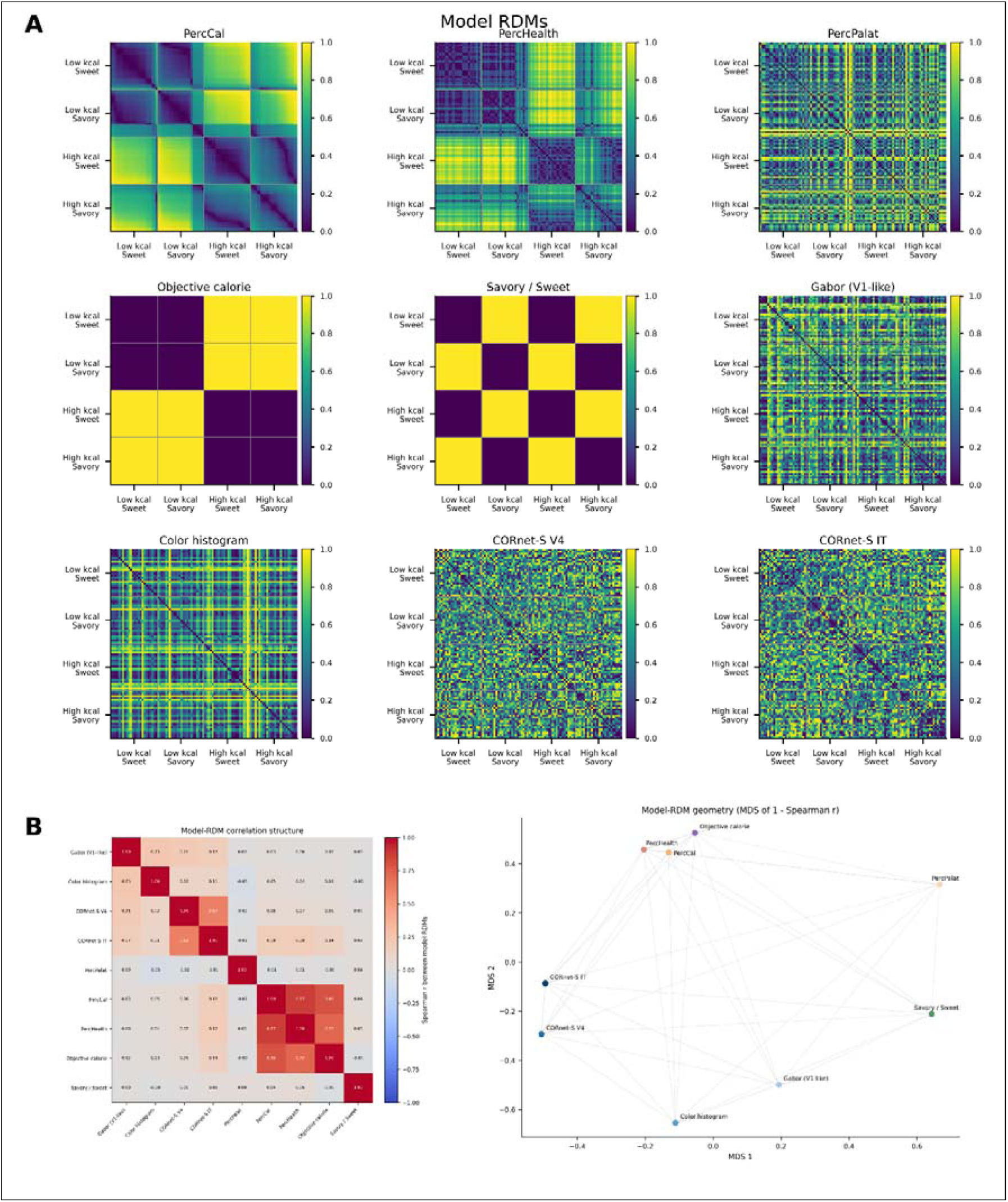
Model RDM structure and relationships among visual, subjective, and categorical food dimensions. (A) Model representational dissimilarity matrices (RDMs) used to characterize the stimulus space. Group subjective RDMs are shown for perceived calorie, perceived health, and perceived palatability. These models were constructed by averaging z-scored ratings across participants for each stimulus and computing pairwise absolute differences between stimuli. Objective categorical RDMs are shown for objective calorie category and savory/sweet category. Visual model RDMs are shown for Gabor-based V1-like features, color histograms, and CORnet-S V4 and IT features. Stimuli are ordered according to the factorial stimulus categories: low-calorie sweet, low-calorie savory, high-calorie sweet, and high-calorie savory. Color values indicate scaled dissimilarity values within each RDM. (B) Model–model representational geometry. Left: Spearman correlations between vectorized model RDMs. Right: two-dimensional multidimensional scaling (MDS) of model distances defined as 1 – ρ (Spearman). The MDS visualization summarizes how similarly different models organize the food stimulus set. The group perceived-calorie and group perceived-health models clustered closely with objective calorie structure, consistent with a shared subjective calorie/health axis, whereas group perceived palatability and the visual models occupied more distinct regions of model space.

### RSA implementation and group inference

All RSA analyses were implemented in Python using custom analysis scripts operating on the fsLR-91k CIFTI beta estimates generated by the first-level GLM. Model–brain correspondence was quantified at the single-subject level for each bilateral Glasser parcel using Spearman rank correlation between vectorized neural and model RDMs. For ROI-level summaries, grayordinates from all parcels belonging to a given ROI group were pooled before neural RDM computation. For each participant and ROI, a single neural RDM was computed from the stimulus-specific response patterns across the pooled grayordinates and correlated with each model RDM. Subject-level model–brain correlations were Fisher-z transformed for group-level averaging and inference, and back-transformed to correlation scale for visualization and reporting.

Group-level inference was performed using a permutation test (i.e. sign-flipping across subjects, 50,000 permutations) on the mean correlation across participants, testing whether model–brain correspondence was reliably greater than zero. To correct for multiple comparisons across models within each ROI, p-values were adjusted using the Benjamini–Hochberg false discovery rate (FDR) procedure. Results were considered significant at an FDR of q<0.05. Unless otherwise stated, q-values reported for ROI-level RSA effects therefore refer to this within-ROI correction family, matching the significance markers shown in the figures. Additional correction summaries across ROIs for each model and across all ROI × model tests are provided in the supplementary tables for transparency.

### Residualized RSA control analyses

As complementary controls, we performed subject-level rank-based residualized RSA. For each subject, the neural RDM vector and target model RDM vector were rank-transformed and separately residualized with respect to a reference model using linear regression; the residualized vectors were then correlated. These analyses tested whether a target model remained related to neural representational geometry after accounting for a specific visual or food-property model. We used this approach to test whether group perceived-calorie and objective-calorie RDMs remained reliable after controlling separately for Color, CORnet V4, CORnet IT, and group perceived health. The visual controls assessed whether calorie-related effects survived image-computable similarity structure, whereas the perceived-health control assessed whether calorie effects reflected separable calorie-related variance or shared calorie/health structure. These residualized RSA analyses were treated as control analyses, not as formal decompositions of unique and shared explained variance, which was instead addressed using the commonality analyses described below.

### Commonality analyses

To formally assess unique and shared contributions of model RDMs, we additionally performed targeted commonality analyses following the logic of Groen et al., 2018. For each participant and ROI, neural RDMs were computed from all valid grayordinates within the pooled ROI and entered into multiple-regression RSA in rank-transformed RDM space. Three theoretically motivated partitions were tested: (1) Visual block + group perceived calorie + group perceived health, to assess whether perceived-calorie or perceived-health structure contributed unique variance beyond their shared subjective calorie/health axis; (2) Visual block + objective calorie + group perceived health, to test whether objective calorie explained unique variance beyond perceived-health structure; and (3) Visual block + objective calorie + group perceived calorie, to test whether objective and perceived calorie contributed separable variance. These partitions were motivated by the high correlations between perceived calorie and perceived health (Spearman ρ = .87), objective calorie and perceived health (ρ = .72), and objective and perceived calorie (ρ = .80).

Across partitions, the Visual block comprised Gabor, Color, CORnet V4, and CORnet IT RDMs. These RDMs were entered as separate rank-transformed predictors within the same block, rather than averaged, so visual commonality components reflect the combined visual model family. This block-wise formulation tested whether calorie- and health-related structure explained neural representational variance beyond a broad set of image-computable similarities.

Commonality inference used stimulus-label permutation. Condition labels of the neural RDM were shuffled, the full decomposition was recomputed for each permutation, and observed components were compared with the resulting null distribution. Unique components were tested with one-sided permutation tests against chance, whereas shared components were tested with two-sided permutation tests because shared terms can be positive or negative. FDR correction was applied across ROIs and components within each partition.

## Results

### Behavioural results

Hunger ratings did not differ significantly between the first (M = 48.36, SD = 15.42) and second session (M = 48.24, SD = 18.36), *t*(24) = 0.05, *p* = 0.95, *d* = 0.01. Likewise, the time elapsed between the last meal and the start of the scanning session (target interval: 120 min) was comparable across sessions (Session 1: M = 131 min, SD = 23.80; Session 2: M = 130.40 min, SD = 21.83), *t*(24) = 0.14, *p* = 0.88, *d* = 0.03. Performance on the orthogonal in-scanner color-discrimination task was high overall (mean run-wise accuracy across participants = 93.4%, SD = 16.5%; mean RT = 776 ms, SD = 120 ms), indicating that participants were generally attentive during stimulus presentation.

### Model RDM structure, behavioral-model collinearity, and reliability checks

Before testing model–brain correspondence, we examined relationships among the model RDMs (Figure 2B). Group perceived calorie was strongly correlated with group perceived health (Spearman ρ = .869), indicating that foods judged as higher in calories were generally judged as less healthy. Perceived calorie was also strongly related to objective calorie category (ρ = .803), and perceived health showed a similarly strong relationship with objective calorie category (ρ = .719). By contrast, perceived palatability and savory/sweet structure were largely independent of the other food-related models (absolute ρ ≤ .049 and ≤ .050, respectively).

Among visual models, CORnet V4 and CORnet IT were most strongly related (ρ = .616), consistent with their shared origin in the same hierarchical visual network. Gabor showed moderate correlations with Color (ρ = .232), CORnet V4 (ρ = .209), and CORnet IT (ρ = .172), but weak relationships with subjective and categorical food models. Perceived calorie and perceived health showed weak-to-moderate correlations with CORnet IT (PercCal: ρ = .176; PercHealth: ρ = .182), and weaker relationships with Color and CORnet V4. Thus, the model space showed substantial collinearity among perceived calorie, perceived health, and objective calorie category, but weaker overlap between these food-related dimensions and the tested visual models. This motivated the residualized RSA controls and commonality analyses reported below.

We also assessed reliability of the group-level behavioral models across rating participants (Supplementary Table S6). Because each participant rated each stimulus once per dimension, within-participant test–retest reliability could not be estimated. Instead, split-half consistency was estimated by repeatedly dividing participants into two balanced halves, computing group-average rating vectors and absolute-difference RDMs separately for each half, and correlating them across 10,000 random splits. Perceived calorie and perceived health were highly reliable as stimulus-wise rating vectors (split-half ρ = .949 and .948; Spearman–Brown corrected ρ = .974 for both) and as model RDMs (split-half ρ = .938 and .942; Spearman–Brown corrected ρ = .968 and .970). Thus, the strong calorie/health relationship reflected stable group-level rating structure. Palatability was less reliable after conversion to an absolute-difference RDM (split-half ρ = .532; Spearman–Brown corrected ρ = .694), consistent with greater idiosyncrasy across participants.

To complement the model-space visualizations, we also inspected neural representational structure directly. Group-average neural RDMs for all predefined ROIs are shown in Supplementary Figure S1, with stimuli ordered according to the same factorial categories used for the model RDMs. These visualizations are descriptive only; all inferential analyses were performed at the subject level. Neural RDMs showed clearer stimulus-linked structure in visual and occipitotemporal ROIs than in OFC, insula, and DLPFC, consistent with the corresponding split-half neural RDM noise ceilings.

### Univariate category-level results

To complement the multivariate representational analyses, we performed a whole-brain univariate contrast comparing responses to high-calorie versus low-calorie food stimuli. Group-level inference was conducted using a random-effects one-sample t-test on subject-level contrast maps, assessing whether category-related activation differences were consistent across participants.

For the high-calorie > low-calorie contrast, significant effects were primarily observed in visual and ventral temporal cortex (Fig. 3B). ROI summaries (Fig. 3C–D) showed reliable positive effects in V1 (mean contrast = 13.42 ± 3.03 SEM, t(24) = 4.43, q = .001, dz = 0.89), VOTC (mean = 6.23 ± 2.02, t(24) = 3.08, q = .010, dz = 0.62), and OFC (mean = 2.05 ± 0.62, t(24) = 3.32, q = .009, dz = 0.66). LOTC did not show a reliable mean activation difference (mean = −2.36 ± 3.48, t(24) = −0.68, q = .605, dz = −0.14), nor did insula (mean = 0.89 ± 0.98, t(24) = 0.91, q = .560, dz = 0.18) or DLPFC (mean = 0.37 ± 1.54, t(24) = 0.24, q = .812, dz = 0.05). Thus, high- versus low-calorie foods elicited reliable mean-response differences mainly in V1, VOTC, and OFC, whereas the RSA analyses tested whether distributed response patterns encoded finer-grained representational structure beyond these mean activation effects.

**Figure 3.**
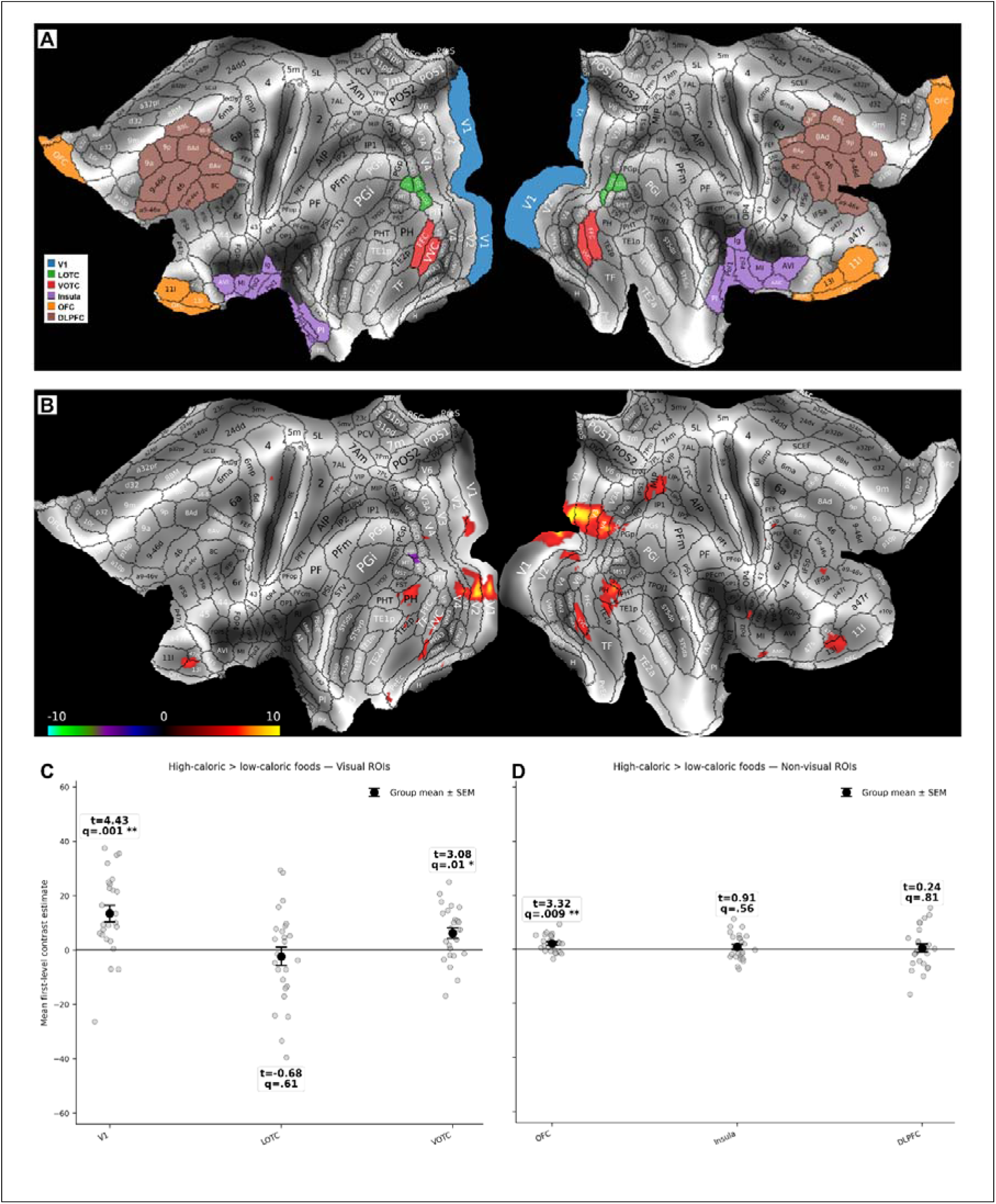
Univariate caloric-category effects and ROI definitions. (A) Predefined regions of interest shown on the HCP-MMP cortical flatmap: V1, lateral occipitotemporal cortex (LOTC), ventral occipitotemporal cortex (VOTC), orbitofrontal cortex (OFC), insula, and dorsolateral prefrontal cortex (DLPFC). (B) Whole-cortex random-effects univariate map for high-caloric > low-caloric foods, shown as thresholded group-level t-values. Warm colors indicate stronger responses to high-caloric foods; cool colors indicate the reverse contrast. (C) ROI-level contrast estimates for visual ROIs. Dots show individual participants, black markers show mean ± SEM, and annotations report t-statistics and FDR-corrected q-values. Significant positive effects were observed in V1 and VOTC, but not LOTC. (D) ROI-level contrast estimates for non-visual ROIs. OFC showed a small positive effect, whereas insula and DLPFC did not survive correction.

The savory > sweet contrast showed a more restricted pattern: after correction, only V1 showed a reliable positive effect (mean = 11.46 ± 3.68 SEM, t(24) = 3.12, q = .028, dz = 0.62), whereas LOTC, VOTC, OFC, insula, and DLPFC did not survive correction.

### Whole-cortex surface parcel-wise RSA

Because the final RSA analyses were performed in fsLR/CIFTI space, we implemented the spatially unbiased analysis as whole-cortex parcel-wise RSA across bilateral Glasser parcels. Of the 180 bilateral HCP-MMP parcels, 179 yielded valid group-level estimates after grayordinate masking. For each model, inference was corrected across parcels using FDR on directional permutation p-values testing positive model–brain correspondence. In the surface maps, colored parcels indicate model-wise FDR-corrected effects, color intensity reflects the mean Spearman correlation across participants, and percentages indicate the mean effect as a percentage of the parcel’s split-half noise ceiling where available. Complete parcel-wise statistics are reported in Supplementary Table S1.

The analysis first confirmed the expected visual-model distribution. Gabor correspondence was confined to early visual cortex, surviving in 17 parcels and peaking in V2 (mean ρ = .366, q < .001), V1 (.355, 49% of NC, q < .001), V3 (.346, q < .001), and V4 (.296, q < .001). CORnet-S V4 and IT showed broader posterior effects, surviving in 34 and 33 parcels, respectively, with peaks in early visual cortex and extension into lateral and ventral occipitotemporal cortex. For CORnet-S IT, corrected effects reached LO1 (.052, 11% of NC, q < .001), LO2 (.046, 11% of NC, q < .001), VVC (.041, 11% of NC, q < .001), LO3 (.025, 6% of NC, q < .001), and FFC (.023, 6% of NC, q < .001). The Color model survived correction in no parcel.

Food-related models showed overlapping posterior occipitotemporal and ventral-temporal distributions. Group perceived calorie survived correction in 18 parcels, strongest in VVC (ρ = .034, 9% of NC, q < .001), PH (.027, q = .002), V8 (.026, q < .001), FST (.025, q = .001), and LO1 (.022, 5% of NC, q < .001). Within the core ventral parcels, the effect was reliable in VVC but not FFC (.010, q = .163). Objective calorie showed a broader pattern, surviving in 30 parcels with similar posterior and ventral-temporal peaks, including VVC (.034, q < .001), PH (.029, q < .001), and V3CD (.027, q < .001), and unlike perceived calorie also survived in FFC (.016, 4% of NC, q < .001). Perceived health survived in 15 parcels with a closely overlapping distribution, strongest in V4t (.038, q < .001), VVC (.035, 9% of NC, q < .001), PH (.030, q < .001), FST (.028, q < .001), and LO1 (.026, 5% of NC, q < .001). Palatability survived in no parcel. Savory/sweet survived weakly in three parcels: LO2 (.011, q = .043), PH (.010, q = .020), and FFC (.007, q = .020). Thus, perceived calorie, objective calorie, and perceived health occupied the same broad posterior occipitotemporal territory, with objective calorie being the most spatially extensive and the only one reliably reaching FFC.

Residualized controls clarified this overlap. After controlling for perceived health, perceived calorie no longer survived correction in occipitotemporal or ventral-temporal cortex; surviving effects were limited to early visual cortex (V2 .042, q = .001; V1 .032, 4% of NC, q = .001; V3 .031, q = .001; V4 .025, q = .004) and a small number of frontal/parietal parcels. The reciprocal control was similarly limited: perceived health controlling for perceived calorie survived only in V4t (.040, q = .029) and MT (.024, q = .041). Given the strong calorie/health collinearity (ρ = .87), and the absence or negativity of the corresponding zero-order effects in early visual cortex, we interpret the early-visual residual effects as a suppression pattern rather than evidence for separable calorie- or health-specific structure in V1/V2. Thus, in the regions where perceived calorie or perceived health were reliable, the two models were not separable, consistent with a shared calorie/health axis.

By contrast, perceived calorie was robust to visual controls. It remained corrected after controlling for Color (15 parcels; VVC .033, 9% of NC, q < .001), CORnet-S V4 (16 parcels; VVC .033, 9% of NC, q = .002), and CORnet-S IT (9 parcels; VVC .028, 7% of NC, q = .002; FST .022, q = .005; PH .021, q = .035; LO1 .013, q = .041). Thus, the occipitotemporal perceived-calorie effect was not reducible to the tested visual models. Objective calorie showed even stronger robustness to visual controls, surviving after Color (29 parcels; FFC .016, q < .001), CORnet-S V4 (27 parcels; FFC .014, q = .001), and CORnet-S IT (19 parcels; FFC .013, q = .003), but again collapsed under perceived-health control, leaving only early-visual suppression effects (e.g., V2 .038, q < .001; V1 .032, q < .001).

Finally, subject-specific behavioral RDMs tested whether each participant’s own rating geometry matched their neural geometry. These analyses converged with the group-level results. Subject-specific perceived calorie survived correction in 13 parcels within the same posterior and ventral-temporal territory, strongest in VVC (.033, 9% of NC, q = .001), V3CD (.030, q = .002), PH (.028, q = .008), and LO1 (.027, 6% of NC, q = .005), and also reached a small corrected effect in FFC (.015, q = .040). Subject-specific perceived health survived in 17 parcels with a highly overlapping distribution, strongest in VVC (.035, 9% of NC, q = .002), V3CD (.028, q = .006), and V4t (.028, q = .002), including FFC (.020, q = .014). Subject-specific palatability survived in no parcel. Thus, individual-rating analyses reproduced reliable perceived-calorie and perceived-health structure in occipitotemporal and ventral-temporal cortex, and the absence of palatability effects, without relying on group-average behavioral models.

### ROI-level summaries of predefined parcel groups

After establishing the spatial distribution of effects in the whole-cortex analysis, we summarized model correspondence within predefined ROI groups motivated by prior literature (Figure 3A). ROI-level effects are shown in Figure 5 and reported in full in Supplementary Table S7. Grayordinates from all Glasser parcels within an ROI were pooled before neural-RDM computation, rather than averaging parcel-level estimates, so these analyses directly tested whether the pooled ROI pattern expressed each model geometry. Unless stated otherwise, q-values refer to FDR correction across models within each ROI. Percentages indicate the mean effect as a fraction of the pooled-ROI split-half noise ceiling.

**Figure 4.**
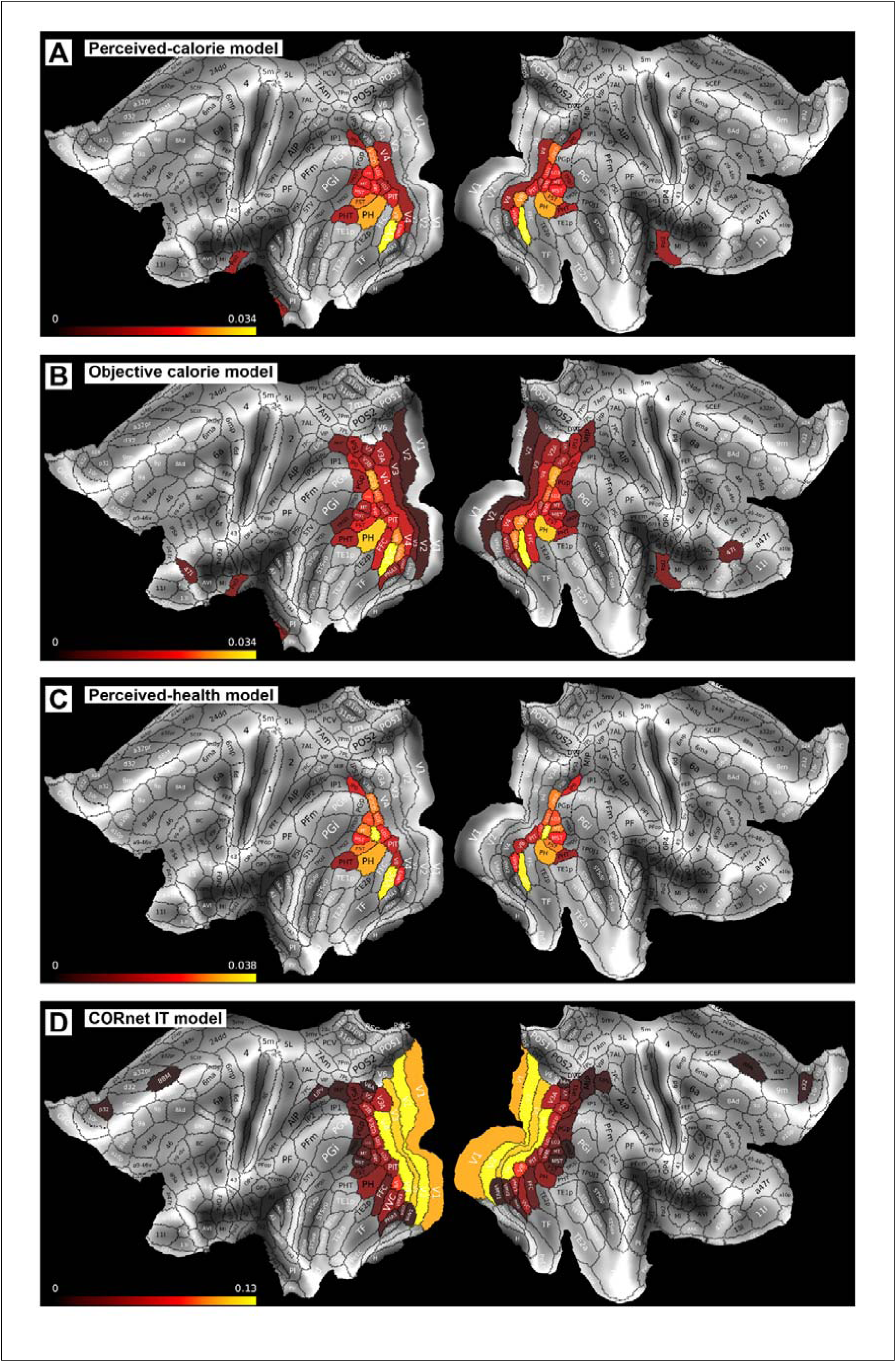
Parcel-wise RSA maps for group perceived-calorie objective-calorie, group perceived-health and visual model RDMs. Whole-cortex bilateral Glasser parcel-wise RSA maps showing group-level model–brain correspondence for A) group perceived calorie, B) objective calorie category, C) group perceived health, and D) CORnet-S IT. Values represent mean Spearman correlations between neural and model RDMs across participants. Colored parcels indicate positive effects surviving model-wise FDR correction across the 179 analyzable bilateral parcels; non-significant parcels are shown in gray. Color scales show thresholded positive effects and are scaled separately for each model. Perceived calorie showed corrected effects mainly in posterior visual, lateral occipitotemporal, ventral temporal, and parahippocampal parcels, strongest in VVC. Objective calorie showed a broader overlapping pattern including VVC, PH, V3CD, V8, VMV3, LO1/LO3, PIT, and FFC. Perceived health showed a similar posterior occipitotemporal distribution, strongest in V4t, VVC, PH, MT, FST, and V3CD. CORnet-S IT showed strongest correspondence in early/intermediate visual cortex, extending into lateral and ventral occipitotemporal parcels.

**Figure 5.**
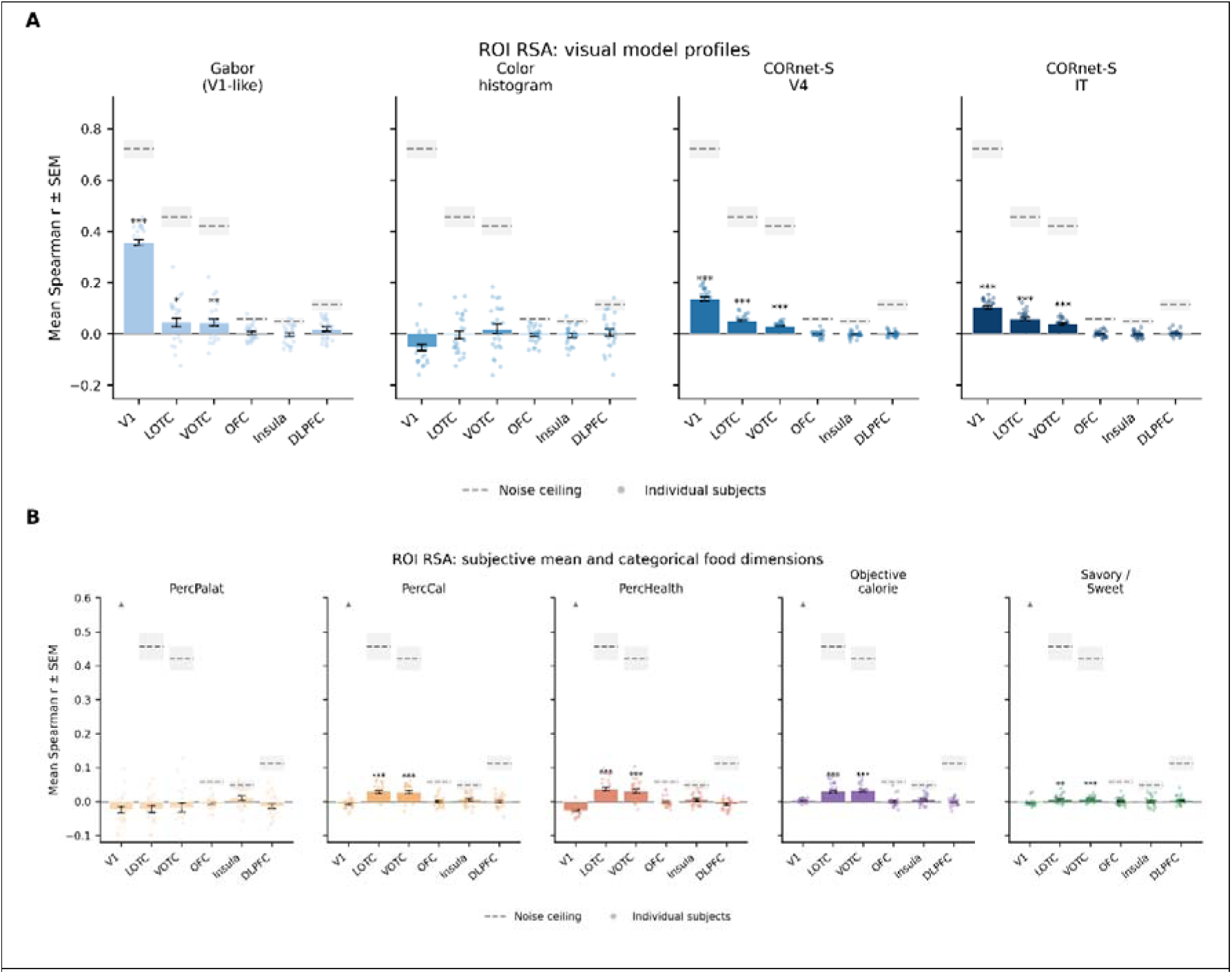
Primary RSA model results for visual and food-related dimensions. A. ROI-level RSA for visual feature models. Models included Gabor, Color, CORnet-S V4, and CORnet-S IT. Gabor showed strongest correspondence in V1, whereas CORnet-S V4 and IT showed reliable positive correspondence in V1, LOTC, and VOTC, consistent with higher-level visual structure in occipitotemporal cortex. Color did not show reliable positive correspondence. B. ROI-level RSA for group subjective and categorical food models: perceived palatability, perceived calorie, perceived health, objective calorie category, and savory/sweet category. Perceived calorie, perceived health, and objective calorie showed reliable positive correspondence in LOTC and VOTC, consistent with shared occipitotemporal food-related structure. Savory/sweet effects were smaller, while palatability showed no reliable positive correspondence and was descriptively negative in visual ROIs. Higher-order ROIs showed no robust positive food-model effects. Bars show group mean Spearman correlations, error bars show ± SEM, dots show individual participants, and gray dashed lines/shaded regions indicate mean neural noise ceiling ± SEM. Asterisks denote effects surviving Benjamini–Hochberg FDR correction across models within each ROI.

In V1, representational structure was dominated by visual information. Gabor was strongest (mean ρ = .355, 49% of NC, q < .001), followed by CORnet-S V4 (.135, 18% of NC, q < .001) and CORnet-S IT (.104, 14% of NC, q < .001); Color was not reliable (−.053, q = 1.000). No subjective food model was reliable (PercCal −.007, q = 1.000; PercHealth −.026, q = 1.000), and objective calorie showed only a negligible positive effect (.003, q = .043). Palatability and SavorySweet were not reliable.

In LOTC, neural geometry corresponded to both visual and food-related models. Visual effects were reliable for Gabor (.044, 9% of NC, q = .010), CORnet-S V4 (.051, 11% of NC, q < .001), and CORnet-S IT (.058, 12% of NC, q < .001). Food-related models were also reliable: PercCal (.028, 6% of NC, q < .001), PercHealth (.037, 8% of NC, q < .001), and objective calorie (.030, 6% of NC, q < .001). SavorySweet was reliable but very small (.006, 1% of NC, q = .008), whereas palatability was descriptively negative and not interpreted as positive correspondence.

In VOTC, pooled ROI analyses showed reliable visual, perceived-calorie, perceived-health, and objective-calorie effects. Visual correspondence was reliable for Gabor (.044, 10% of NC, q = .002), CORnet-S V4 (.030, 7% of NC, q < .001), and CORnet-S IT (.039, 9% of NC, q < .001), while Color was positive but not corrected (.018, q = .185). The three correlated food-related models were all reliable: PercCal (.028, 6% of NC, q < .001), PercHealth (.030, 7% of NC, q < .001), and objective calorie (.032, 7% of NC, q < .001). SavorySweet was also weakly reliable (.007, 1% of NC, q < .001), and palatability was descriptively negative. Thus, VOTC showed positive correspondence with perceived calorie, perceived health, and objective calorie category, but the control analyses below indicate that these effects largely reflected shared rather than separable structure.

As a descriptive check, we visualized group-average neural geometry in occipitotemporal ROIs using MDS plots with individual food images overlaid (Supplementary Figure S2). These plots provide an intuitive view of stimulus organization in ROI-level neural response-pattern space but were not used for inference. Linear gradients of stimulus properties were fitted across the two-dimensional MDS solutions to aid interpretation; the resulting MDS-gradient R² values summarize visualization structure only and should not be interpreted as neural variance explained or RSA effect sizes.

Higher-order ROIs showed no robust positive food-model effects, and pooled-ROI RDM reliability was near zero. In OFC, no model survived correction (PercCal −.0001, q = .819; PercHealth −.002, q = .819; objective calorie .000, q = .819; SavorySweet .001, q = .819). In insula, none survived correction, although several effects were nominally positive before correction (PercCal .005, p = .087; PercHealth .006, p = .078; palatability .010, p = .047; objective calorie .005, p = .053; all q = .148). In DLPFC, no model survived correction (PercCal .001, q = .975; PercHealth −.009, q = .998; objective calorie −.001, q = .998; SavorySweet .003, q = .830). Overall, the higher-order ROIs provided no robust evidence for the tested food-rating geometries under the present approach.

Residualized RSA controls (Figure 6; full values in Supplementary Table S7) showed that LOTC and VOTC perceived-calorie effects were robust to visual controls but not separable from perceived-health structure. In LOTC, PercCal remained reliable after controlling for Gabor (ρ = .028, 6% of NC, SEM = .005, q < .001), Color (.029, 6% of NC, SEM = .005, q < .001), CORnet V4 (.026, 5% of NC, SEM = .005, q < .001), and CORnet IT (.019, 4% of NC, SEM = .004, q < .001). However, it was no longer reliable after controlling for PercHealth (−.007, SEM = .008, q = .831). The reciprocal control showed a reduced but reliable PercHealth effect after controlling for PercCal (.024, 5% of NC, SEM = .009, q = .008). We do not interpret this as evidence for a separable health-specific code; rather, the LOTC perceived-calorie effect appears robust to visual controls but not specific to calorie, and is better interpreted as shared calorie/health structure.

**Figure 6.**
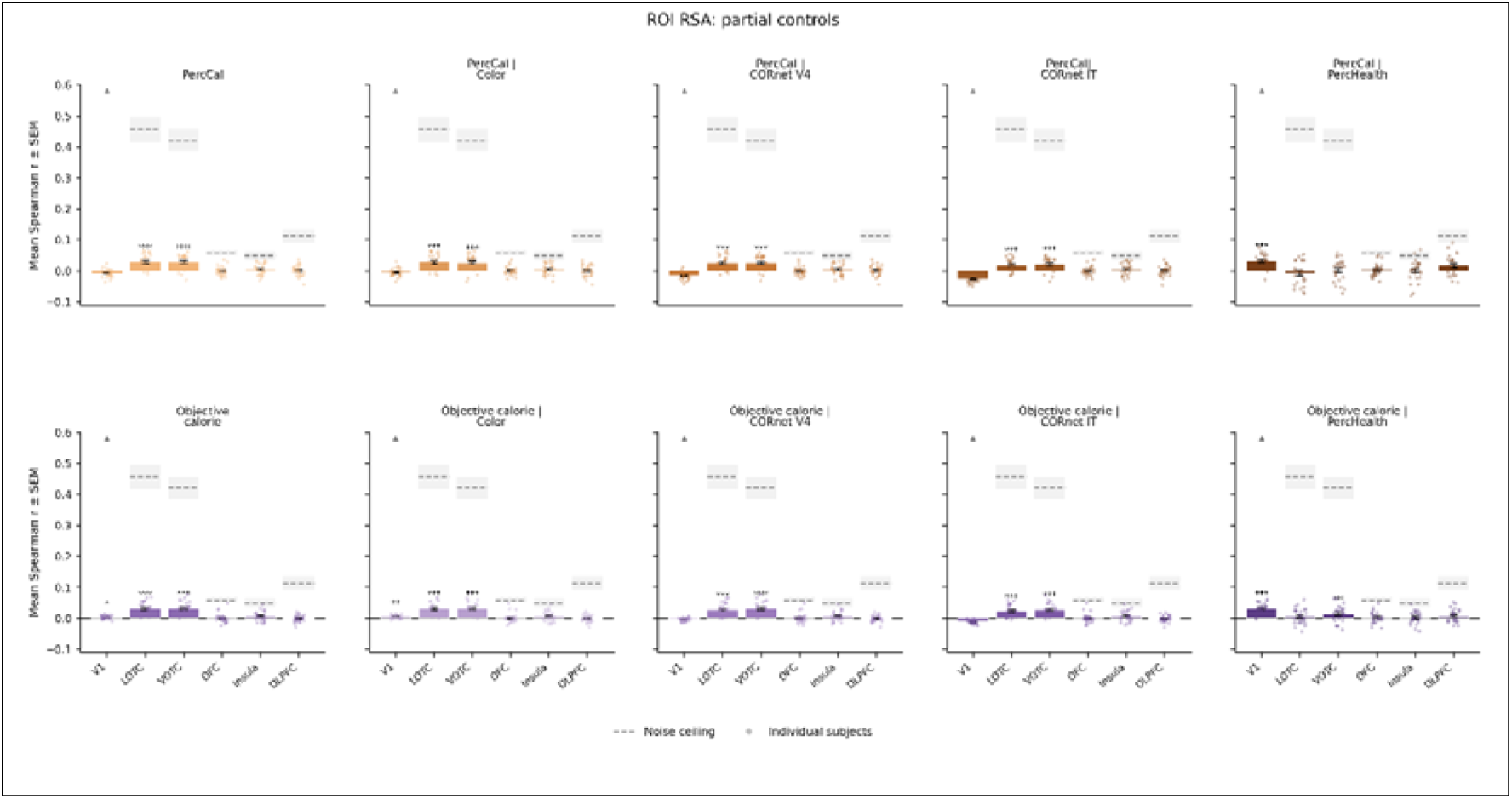
Partial RSA controls for group perceived-calorie and objective-calorie models. Partial RSA tested whether calorie-related effects were explained by visual similarity or overlap with perceived-health structure. The top row shows group perceived calorie before and after controlling for Color, CORnet-S V4, CORnet-S IT, and perceived health; the bottom row shows objective calorie category with the same controls. Perceived-calorie effects remained reliable in LOTC and VOTC after visual controls, but not after controlling for perceived health, indicating shared subjective calorie/health structure rather than a separable perceived-calorie representation. Objective calorie also remained reliable in LOTC and VOTC after visual controls, indicating a small categorical calorie effect beyond the tested visual models; after perceived-health control, this effect was attenuated and remained reliable only in VOTC. Bars show group mean Spearman correlations, error bars show ± SEM, dots show individual participants, and gray dashed lines/shaded regions indicate mean neural noise ceiling ± SEM. Asterisks denote effects surviving Benjamini–Hochberg FDR correction across models within each ROI.

VOTC showed the same pattern. PercCal remained reliable after controlling for Gabor (.026, 6% of NC, SEM = .005, q < .001), Color (.027, 6% of NC, SEM = .005, q < .001), CORnet V4 (.026, 5% of NC, SEM = .005, q < .001), and CORnet IT (.021, 4% of NC, SEM = .004, q < .001), but not after controlling for PercHealth (.002, SEM = .007, q = .386). The reciprocal control was also non-significant (PercHealth controlling for PercCal: .013, SEM = .008, q = .074), indicating symmetric non-separability between calorie and health in VOTC.

Objective calorie category was more robust to visual controls. In LOTC, CalorieObjective remained reliable after controlling for Gabor (.030, 6% of NC, SEM = .004, q < .001), Color (.031, 6% of NC, SEM = .004, q < .001), CORnet V4 (.028, 6% of NC, SEM = .004, q < .001), and CORnet IT (.022, 5% of NC, SEM = .004, q < .001), but not after controlling for PercHealth (.006, SEM = .005, q = .146). In VOTC, CalorieObjective remained reliable after all visual controls (Gabor: .031, 6% of NC, SEM = .004, q < .001; Color: .030, 7% of NC, SEM = .004, q < .001; CORnet V4: .030, 6% of NC, SEM = .004, q < .001; CORnet IT: .026, 6% of NC, SEM = .004, q < .001) and also remained reliable, though reduced, after controlling for PercHealth (.014, 3% of NC, SEM = .005, q = .009). Thus, objective calorie contributed to occipitotemporal representational structure beyond the tested visual models, but its attenuation under perceived-health control indicates strong overlap with the broader calorie/health structure of the stimulus set.

Because perceived calorie and perceived health were highly collinear (ρ = .87), these partial controls were used to assess non-separability rather than to claim unique codes. The null calorie-controlling-for-health results indicate that calorie structure was not separable from health in occipitotemporal ROIs. Conversely, positive partial effects between collinear models were not interpreted as evidence for distinct codes; formal partitioning of unique and shared variance was addressed by the commonality analysis. Consistent with a suppression interpretation, health-residualized controls produced small positive effects in V1, where the corresponding zero-order calorie and health effects were null or negative: PercCal controlling for PercHealth (.032, 4% of NC, SEM = .005, q < .001) and CalorieObjective controlling for PercHealth (.032, 4% of NC, SEM = .003, q < .001). We therefore interpret these early-visual residual effects as arising from calorie/health collinearity rather than as evidence for abstract calorie coding in V1.

As an additional subject-specific RSA, we tested whether each participant’s own subjective rating geometry predicted their neural representational geometry. For each participant and rating dimension, ratings were z-scored across stimuli and converted into pairwise absolute-difference RDMs. These subject-own models were analyzed separately from the shared group-level RDMs because the model vector differed across participants. Full subject-own ROI statistics are reported in Supplementary Table S5; percentages indicate the mean RSA effect as a fraction of the corresponding ROI’s r-space neural RDM noise ceiling.

At the ROI level, subject-own perceived-calorie and perceived-health models showed reliable positive effects in both LOTC and VOTC. In LOTC, subject-own perceived calorie was reliable (mean ρ = .029, 6% of NC, SEM = .007, p < .001, q < .001), as was subject-own perceived health (ρ = .032, 6% of NC, SEM = .005, p < .001, q < .001). Subject-own palatability was descriptively negative and not positively reliable (ρ = −.018, −4% of NC, SEM = .009, q = .966). In VOTC, subject-own perceived calorie was similarly reliable (ρ = .030, 6% of NC, SEM = .006, p < .001, q < .001), as was subject-own perceived health (ρ = .034, 7% of NC, SEM = .006, p < .001, q < .001), whereas palatability was again descriptively negative and not reliable (ρ = −.022, −5% of NC, SEM = .010, q = .985). No subject-own model survived correction in V1, OFC, insula, or DLPFC.

Overall, subject-own analyses indicate that participant-specific perceived-calorie and perceived-health geometry is reflected in occipitotemporal neural representational structure, with convergent ROI-level effects in LOTC and VOTC and parcel-wise effects centered on VVC, LO/V3CD, PH, FST, and related posterior occipitotemporal parcels. In contrast, subject-specific palatability geometry showed no reliable positive ROI-level or parcel-wise correspondence.

### Commonality analysis

To formally evaluate unique and shared model contributions, we performed targeted commonality analyses using multiple-regression RSA in rank-transformed RDM space. These analyses decomposed the variance explained by theoretically motivated model sets into unique and shared components. Because commonality coefficients can be difficult to interpret in sign when predictors are strongly correlated, we focus primarily on unique variance estimates and treat reliable shared terms as evidence for overlapping representational structure.

In the first partition, the Visual block was entered together with group perceived calorie and group perceived health. This tested whether either subjective model explained neural representational variance beyond visual similarity and beyond their shared subjective calorie/health structure, given their strong correlation (Spearman ρ = 0.87). The full model explained the most variance in V1 (R² = 16.23%), followed by LOTC (R² = 2.23%) and VOTC (R² = 1.91%). The unique Visual component was reliable in V1 (16.00% R², SEM = 0.92%, p < .001, q = .006) and LOTC (1.86% R², SEM = 0.30%, p = .002, q = .010), but not after correction in VOTC (1.61% R², SEM = 0.29%, p = .021, q = .098). Critically, neither perceived calorie nor perceived health explained reliable unique variance in LOTC or VOTC. In LOTC, unique perceived calorie (0.15% R², SEM = 0.03%, p = .509, q = .652) and unique perceived health (0.22% R², SEM = 0.05%, p = .114, q = .281) were both non-significant. The same was true in VOTC for perceived calorie (0.13% R², SEM = 0.03%, p = .792, q = .876) and perceived health (0.18% R², SEM = 0.05%, p = .512, q = .652). However, the shared Visual × perceived-calorie × perceived-health component was reliable in both LOTC and VOTC (both 0.04% R², SEM = 0.01%, p < .001, q = .006), indicating overlapping rather than separable subjective calorie/health-related structure.

In the second partition, the Visual block was entered together with objective calorie and group perceived health. This tested whether objective calorie category explained unique variance beyond perceived-health structure, given their strong correlation (Spearman ρ = 0.72). The full model explained 16.22% of neural-RDM variance in V1, 2.15% in LOTC, and 1.87% in VOTC. The unique Visual component was reliable in V1 (16.01% R², SEM = 0.92%, p < .001, q = .005) and LOTC (1.87% R², SEM = 0.31%, p = .002, q = .008), but not after correction in VOTC (1.61% R², SEM = 0.29%, p = .024, q = .077). Neither objective calorie nor perceived health showed reliable unique variance in LOTC or VOTC. In LOTC, unique objective calorie (0.06% R², SEM = 0.02%, p = .211, q = .368) and unique perceived health (0.13% R², SEM = 0.04%, p = .083, q = .183) were non-significant. In VOTC, unique objective calorie was positive but did not survive correction (0.09% R², SEM = 0.02%, p = .029, q = .084), and unique perceived health was also non-significant (0.12% R², SEM = 0.03%, p = .255, q = .396). By contrast, shared components involving objective calorie and perceived health were reliable in both LOTC and VOTC, although small in magnitude. In LOTC, both the shared objective-calorie × perceived-health component (0.03% R², SEM = 0.02%, p < .001, q = .005) and the shared Visual × objective-calorie × perceived-health component (0.04% R², SEM = 0.01%, p < .001, q = .005) were reliable. The same pattern was observed in VOTC, where the shared objective-calorie × perceived-health component (0.02% R², SEM = 0.02%, p < .001, q = .005) and the shared Visual × objective-calorie × perceived-health component (0.03% R², SEM = 0.01%, p < .001, q = .005) were reliable. These results support overlapping objective-calorie/perceived-health structure rather than separable unique effects.

In the third partition, the Visual block was entered together with objective calorie and group perceived calorie. This tested whether objective and perceived caloric structure contributed separable variance, given their high collinearity (Spearman ρ = 0.80). The full model explained 16.12% of neural-RDM variance in V1, 2.08% in LOTC, and 1.81% in VOTC. The unique Visual component was reliable in V1 (16.04% R², SEM = 0.92%, p < .001, q = .008) and LOTC (1.88% R², SEM = 0.30%, p = .002, q = .014), but not after correction in VOTC (1.61% R², SEM = 0.29%, p = .023, q = .107). Neither calorie model carried reliable unique variance beyond the other in LOTC or VOTC. In LOTC, unique objective calorie (0.06% R², SEM = 0.02%, p = .428, q = .619) and unique perceived calorie (0.06% R², SEM = 0.02%, p = .906, q = .992) were both non-significant. The same was observed in VOTC for objective calorie (0.07% R², SEM = 0.02%, p = .416, q = .619) and perceived calorie (0.06% R², SEM = 0.01%, p = .964, q = .992). However, shared components involving objective and perceived calorie were reliable in both LOTC and VOTC. In LOTC, both the shared objective-calorie × perceived-calorie component (0.03% R², SEM = 0.03%, p < .001, q = .008) and the shared Visual × objective-calorie × perceived-calorie component (0.04% R², SEM = 0.01%, p < .001, q = .008) were reliable. The same was true in VOTC for the shared objective-calorie × perceived-calorie component (0.04% R², SEM = 0.02%, p < .001, q = .008) and the shared Visual × objective-calorie × perceived-calorie component (0.03% R², SEM = 0.01%, p < .001, q = .008). Thus, objective and perceived calorie contributed primarily through shared calorie-related structure rather than separable unique components.

Overall, the commonality analyses support three conclusions. First, visual model structure dominated V1 and explained the largest fraction of neural-RDM variance overall. Second, LOTC and VOTC contained weak but reliable food-related representational structure, but this structure was largely shared across correlated food dimensions. Third, perceived calorie did not explain reliable unique variance beyond perceived health or objective calorie in LOTC or VOTC. Thus, the mean-model results are most consistent with a shared occipitotemporal food-related axis on which perceived calorie, perceived health, and objective calorie partly covary, rather than with a uniquely isolated perceived-calorie code.

## Discussion

The present study investigated how visual and higher-order food properties, including subjective ratings and objective category structure, are encoded across the cortical processing hierarchy. Stimulus-level fMRI analyses showed that food representations were organized along multiple, partly overlapping axes. Visual model structure was strongest in early visual cortex and extended into lateral and ventral occipitotemporal regions. Food-related models showed smaller but reliable effects in occipitotemporal cortex: group perceived calorie, group perceived health, and objective calorie were all associated with neural representational geometry in LOTC and VOTC.

Control analyses indicate that these effects should not be interpreted as an isolated scalar calorie code. Perceived-calorie effects in LOTC and VOTC remained reliable after controlling for visual feature models, but not after controlling for perceived health. Objective calorie also remained reliable after visual controls, but was attenuated after perceived-health control. Commonality analyses converged with this pattern: perceived calorie, perceived health, and objective calorie did not explain robust separable unique variance beyond one another in occipitotemporal cortex, but contributed mainly through shared components. Thus, food-related information appears to be embedded within visual representational spaces in a graded and model-dependent manner, with LOTC and VOTC expressing a shared food-property axis on which perceived calorie, perceived health, and objective calorie partly covary. These findings are based on group-level model RDMs in a sample of healthy young women, a scope we return to in the Limitations.

### Food-related representational structure in occipitotemporal cortex

Contemporary accounts characterize occipitotemporal cortex as a continuous, multidimensional space shaped by visual, semantic, and behaviorally relevant object properties rather than by low-level similarity or discrete category membership alone (Bracci & Peelen, 2013; Bracci, Ritchie, & de Beeck, 2017; DiCarlo et al., 2012; Grill-Spector & Weiner, 2014; Huth et al., 2012; Op de Beeck et al., 2008; Wurm, Ariani, Greenlee, & Lingnau, 2016; Wurm, Caramazza, & Lingnau, 2017). Within this framework, LOTC has been linked to abstract and action-relevant object dimensions that cut across categories (Cortinovis, Peelen, & Bracci, 2025; Kabulska, Zhuang, & Lingnau, 2024; Perini, Caramazza, & Peelen, 2014; Wurm & Lingnau, 2015), whereas VOTC and fusiform-adjacent cortex have been associated with high-level visual categorization, domain selectivity, and stable object representations within ventral temporal cortex (Grill-Spector & Weiner, 2014). Recent work on food perception further suggests that ventral visual and occipitotemporal food representations may reflect not only visual appearance, but also healthfulness, processing level, learned nutritional associations, graspability, and other consumption-related properties (Avery et al., 2025; Carrington et al., 2024; Jain et al., 2023; Khosla et al., 2022; Moerel et al., 2024; Pennock et al., 2023; Ritchie et al., 2024). We therefore asked whether LOTC and VOTC carry food-related structure beyond the tested visual models, using three strongly correlated dimensions (perceived calorie and perceived health ρ = 0.87, perceived calorie and objective calorie ρ = 0.80, objective calorie and perceived health ρ = 0.72) and testing, through residualized RSA and commonality analysis, whether their neural correspondence is separable or shared.

In LOTC, all three food models showed positive effects (perceived calorie 0.028, perceived health 0.037, objective calorie 0.030, all q < .001) that survived the visual controls but not perceived-health control (perceived calorie | perceived health: −0.007, q = .831; objective calorie | perceived health: 0.006, q = .146). The commonality analysis converged: neither perceived calorie nor perceived health explained reliable unique variance beyond the other and the Visual block (unique perceived calorie 0.15%, q = .652; unique perceived health 0.22%, q = .281), whereas their shared visual, calorie, and health component was reliable (0.04%, q = .006), and objective and perceived calorie likewise carried no reliable unique variance beyond one another. LOTC therefore expressed a shared food-property axis rather than separable calorie- and health-specific codes. This did not depend on pooling: perceived calorie survived parcel-wise correction within LO1 (0.022, q < .001), LO2 (0.014, q = .014), and LO3 (0.019, q = .007), and held despite no reliable univariate high- versus low-calorie difference in LOTC (t(24) = −0.68, q = .605), indicating stimulus-level pattern geometry rather than a mean-amplitude effect.

VOTC showed a convergent pattern. The high- versus low-calorie univariate contrast was reliable, consistent with stronger ventral responses to high-calorie foods (Dąbkowska-Mika et al., 2023; Killgore et al., 2003; Siep et al., 2009; Toepel, Knebel, Hudry, le Coutre, & Murray, 2009; Yang, Wu, & Morys, 2021) and all three food models were reliable in RSA (perceived calorie 0.028, perceived health 0.030, objective calorie 0.032, all q < .001). As in LOTC, perceived calorie survived the visual controls but not perceived-health control (0.002, q = .386), while objective calorie retained a small effect after perceived-health control (0.014, q = .009). The commonality analysis, however, found no reliable unique food component: in every partition the unique terms failed correction, while the shared visual and food components were reliable. The VOTC effect is therefore best read as overlapping visual and food-property structure, not a separable calorie-specific representation.

The parcel-wise maps provided a more spatially specific view of this occipitotemporal effect. For group perceived calorie, corrected effects were distributed across posterior visual, lateral occipitotemporal, ventral-temporal, and parahippocampal parcels, with the strongest corrected effect in VVC (mean ρ = 0.034, q < .001) and additional effects in PH (mean ρ = 0.027, q = .002), V8, FST, V3CD, LO1/LO2/LO3, VMV3, MST, V4t, PIT, and related parcels. Notably, perceived calorie did not survive correction in FFC. Group perceived health showed a closely related distribution, again including VVC (mean ρ = 0.035, q < .001) and PH (mean ρ = 0.030, q < .001), but not FFC. Objective calorie showed a broader overlapping pattern: it survived correction in VVC (mean ρ = 0.034, q < .001), PH (mean ρ = 0.029, q < .001), and FFC (mean ρ = 0.016, q < .001), as well as V3CD, V8, VMV3, LO1/LO3, V4t, FST, PIT, V4, IP0, LO2, and additional posterior visual parcels. Thus, within ventral occipitotemporal cortex, VVC showed consistent correspondence with perceived calorie, perceived health, and objective calorie, whereas at the group level FFC showed a corrected effect only for objective calorie.

Taken together, LOTC and VOTC differ in emphasis rather than dissociating. Both carried food-related structure beyond the visual models, with perceived calorie closely tied to perceived health. LOTC showed the subjective calorie/health profile without univariate calorie-category activation, whereas VOTC added univariate calorie sensitivity and a small objective-calorie residual that survived pairwise perceived-health control (q = .009) but not the commonality unique-variance test (q = .084). What neither region provided was evidence for a separable calorie-, health-, or category-specific code. Instead, both expressed a single shared food-property axis, on which perceived calorie, perceived health, objective calorie, and learned food knowledge covary rather than dissociate.

This shared-axis interpretation also fits with recent work arguing that food selectivity in occipitotemporal cortex should not be understood as a discrete food-category response in isolation. Ritchie et al. showed that responses to food can overlap with responses to tools and interpreted this overlap as evidence that food representations are partly embedded within broader object-representation systems organized by behaviorally relevant properties (Ritchie et al., 2024). The present findings extend this perspective from food-versus-object category contrasts to stimulus-level representational geometry within foods. Rather than isolating a single food attribute, such as calorie content, LOTC and VOTC expressed representational structure along a covarying food-property axis spanning visual appearance, perceived calorie, perceived health, objective calorie category, and learned food knowledge. In this sense, our results are consistent with a view of occipitotemporal food representations as multidimensional and behaviorally grounded, rather than as reflecting either a purely visual food category or a narrowly calorie-specific code.

This multidimensional interpretation also aligns closely with prior work identifying a dominant food-related representational dimension organized around healthfulness and processing level in ventral temporal/parahippocampal cortex (Avery et al., 2025). The present results converge with that account in two ways. First, at the model level, our perceived-calorie, perceived-health, and objective-calorie RDMs were strongly correlated, indicating that subjective calorie estimates, health-related judgments, and objective calorie category were not independent dimensions in this stimulus set. Second, at the neural level, the parcel-wise maps showed consistent food-related effects in ventral-temporal and parahippocampal territory, including VVC and PH, with PH providing an especially direct spatial parallel to the region emphasized by Avery et al. An important extension is that this axis was expressed during incidental food viewing, with calorie, health, and processing judgments never task-relevant during scanning. Occipitotemporal cortex therefore encodes a broad food-property axis overlapping the healthfulness/processing dimension Avery et al. described, without that structure depending on explicit evaluation.

This interpretation also connects naturally to visually grounded accounts of food selectivity in ventral visual cortex. Food-selective responses have been associated with chromatic and other low-level visual properties, including the overrepresentation of color-biased regions within food-selective cortex (Henderson et al., 2025; Pennock et al., 2023). Our findings refine rather than contradict this view. The food-related effects persisted after controlling for the tested visual models, but these controls cannot exclude the possibility that other visual regularities, not captured by the present model set, contribute to the observed structure. At the same time, the strong overlap between perceived calorie, perceived health, objective calorie, processing-related food structure, and learned food knowledge indicates that the effect should not be interpreted as purely visual or purely semantic. The present design cannot determine how much of this shared axis is driven by image-computable visual structure versus broader learned food knowledge, because processing level, preparation state, perceived calorie, perceived health, and visual appearance naturally covary across food images. Instead, the results point to a multidimensional food-property axis at the intersection of visual appearance, learned food knowledge, and subjective food evaluation.

A remaining question is what kind of information gives rise to this shared food-property axis. One possibility is that the axis is largely driven by visual appearance: foods that look processed, prepared, dessert-like, or energy-dense also tend to be judged as higher in calorie and lower in health, whereas raw or minimally processed foods tend to occupy the opposite end of the space. Another possibility is that the same axis also reflects learned food knowledge, such as expectations about energy density, ingredients, or preparation, that is only partly available from appearance. The present RSA results show that perceived calorie, perceived health, and objective calorie covary in occipitotemporal representational geometry, but they do not determine why they covary. Thus, the next question is not simply whether calorie, health, and food category are independently represented, but what kinds of visual, semantic, and learned food information jointly define this shared axis. Ultimately, fully separating these possibilities will require stimulus sets and model comparisons designed to decorrelate visual appearance, processing level, perceived health, and caloric density.

### Context dependence of higher-order food representations

Despite strong theoretical motivation, we did not observe robust positive representational correspondence between subjective food-judgment models and neural geometry in OFC, insula, or DLPFC under the present task conditions. This absence of corrected positive RSA effects should not be interpreted as evidence that these regions are generally uninvolved in food valuation. Rather, during an orthogonal fixation-color task and using offline behavioral RDMs, these higher-order regions did not show stable stimulus-level representational structure for perceived calorie, perceived health, or palatability comparable to the effects observed in occipitotemporal cortex. Subject-specific behavioral RDMs led to the same broad conclusion: unlike LOTC and VOTC, OFC, insula, and DLPFC did not show corrected positive effects for participant-specific ratings.

The higher-order ROI results were regionally heterogeneous. In OFC, the whole-brain univariate analysis revealed category-related activation differences for the high-calorie versus low-calorie contrast (Figure 3), indicating sensitivity to caloric category at the level of mean response amplitude. However, this sensitivity did not translate into reliable stimulus-level representational structure in RSA. Bayesian sensitivity analyses, performed with the Bayes Factor toolbox for MATLAB (Krekelberg, 2024; Rouder, Speckman, Sun, Morey, & Iverson, 2009), supported this interpretation: OFC showed moderate evidence for the null for group perceived calorie (mean ρ = −.0001 ± .0029, BF01 = 4.74), group perceived health (ρ = −.0018 ± .0030, BF01 = 4.03), and perceived calorie controlling for perceived health (ρ = .0030 ± .0042, BF01 = 3.80). Subject-own models in OFC showed the same pattern for own perceived calorie (BF01 = 3.74), own perceived health (BF01 = 4.73), and own perceived palatability (BF01 = 4.08). Thus, OFC showed univariate calorie-category sensitivity, but no evidence for a stable positive representational geometry aligned with the tested subjective food-rating models.

In insula, there was no reliable univariate or corrected RSA evidence for food-related structure. Bayesian evidence for the group subjective models was mostly inconclusive: perceived calorie (ρ = .0051 ± .0037, BF01 = 2.03), perceived health (ρ = .0055 ± .0038, BF01 = 1.85), and palatability (ρ = .0105 ± .0060, BF01 = 1.26) did not provide clear support for either the null or a non-zero effect. The residualized perceived-calorie contrast showed moderate evidence for the null (ρ = .0006 ± .0070, BF01 = 4.73), as did the subject-own perceived-calorie model (BF01 = 4.41). Overall, the insula results are best described as weak and inconclusive rather than as strong evidence for either presence or absence of food-related representational geometry.

DLPFC showed a somewhat different but still non-robust pattern. The unresidualized group perceived-calorie model provided moderate evidence for the null (ρ = .0008 ± .0035, BF01 = 4.64), as did subject-own perceived calorie (ρ = .0044 ± .0048, BF01 = 3.25). Group perceived health showed moderate evidence for a negative association (ρ = −.0089 ± .0031, BF10 = 5.42), and perceived calorie controlling for perceived health showed moderate evidence for a positive effect (ρ = .0172 ± .0067, BF10 = 3.09). Because perceived calorie and perceived health were highly collinear, this residualized contrast should be interpreted cautiously. As with the early-visual residualized effects, it may reflect suppression arising from calorie/health collinearity rather than a genuine calorie-specific representation, although an opponent or residual food-evaluative dimension cannot be excluded. Importantly, this DLPFC pattern did not generalize to the unresidualized perceived-calorie model or to participant-specific calorie ratings. We therefore treat it as suggestive and hypothesis-generating, not as evidence for a robust stimulus-level DLPFC calorie code.

A likely explanation is that the present design was more sensitive to stable stimulus-linked representational structure than to context-dependent food-value signals. Representations in OFC, insula, and DLPFC may be less stable and less stimulus-bound than those in occipitotemporal cortex, because food-value signals in these regions depend strongly on hunger, satiety, motivational state, attentional focus, and current task goals (Franssen et al., 2020; Kochs et al., 2023; Piech et al., 2009; Pimpini et al., 2022; Rolls, 2015; Rolls, Feng, Cheng, & Feng, 2023; Siep et al., 2009; Wright et al., 2016). This may also explain why OFC showed category-level mean-response sensitivity but no reliable RSA effect: OFC may distinguish high- from low-calorie foods at the level of average activation without preserving a stable fine-grained geometry aligned with perceived calorie or perceived health.

The present design was optimized to isolate stable stimulus-linked representations: participants performed an orthogonal color-discrimination task, metabolic state was controlled, and responses were averaged across repeated presentations. These choices increased sensitivity to consistent visual and visual-semantic structure, but may have reduced sensitivity to valuation, interoceptive, choice-related, or regulatory signals that depend on explicit evaluation or current goals. This interpretation is consistent with work showing stronger limbic and prefrontal involvement when food representations are probed using naturalistic similarity judgments or explicit pleasantness and self-control tasks (Avery et al., 2025). Thus, the weak and mixed higher-order ROI results should not be read as contradicting prior evidence for food valuation and control signals in OFC, insula, or DLPFC. Rather, they suggest that these regions may encode food value in a more context-dependent format, whereas the present RSA analyses were most sensitive to stable stimulus-linked structure in occipitotemporal and parahippocampal cortex.

### Palatability-related representational effects

Perceived palatability showed a different pattern from the calorie- and health-related models. In the primary group-level RSA, which used directional tests for positive model–brain correspondence, group perceived palatability showed no positive parcel-wise effects surviving model-wise correction. At the ROI level, palatability was descriptively negative in visual regions, including V1 (mean ρ = −.024, SEM = .009), LOTC (ρ = −.022, SEM = .009), and VOTC (ρ = −.018, SEM = .012). Similar negative or near-zero effects were observed in OFC (ρ = −.005, SEM = .004) and DLPFC (ρ = −.014, SEM = .007). Insula showed a small nominally positive effect (ρ = .010, SEM = .006, p = .047), but this did not survive correction within the ROI model family (q = .149). Thus, the group palatability RDM did not show robust positive correspondence with neural representational geometry in either visual or higher-order ROIs. Because the primary tests were directional for positive correspondence, negative values are reported descriptively and are not interpreted as evidence for an inverse palatability code.

This pattern was not explained by the use of group-average palatability ratings. When each participant’s own palatability ratings were used to construct subject-specific RDMs, no ROI showed a corrected positive effect. Subject-own palatability remained descriptively negative in V1 (ρ = −.021, SEM = .010), LOTC (ρ = −.018, SEM = .009), and VOTC (ρ = −.022, SEM = .010), and was negative or near zero in OFC (ρ = −.003, SEM = .005), insula (ρ = .006, SEM = .006), and DLPFC (ρ = −.011, SEM = .007). The parcel-wise subject-own analysis led to the same conclusion: no parcel showed a positive subject-own palatability effect surviving model-wise correction.

Together, these results indicate that palatability did not behave like the shared calorie/health-related structure in the present analyses. Perceived calorie and perceived health showed reliable positive occipitotemporal correspondence, whereas palatability showed no corrected positive effects and was descriptively negative in visual ROIs. This does not imply that palatability is not represented in the brain. Rather, palatability may be more state-dependent, preference-specific, and task-sensitive than the shared calorie/health structure observed here, and may require explicit liking, taste imagery, choice, or consumption-related task demands to be robustly expressed. This interpretation is consistent with evidence implicating insular and mesocorticolimbic regions in gustatory, interoceptive, and hedonic food processing (Craig, 2009; Simmons et al., 2013; Small, 2010), and with work showing that taste- or palatability-focused processing can preferentially engage these systems (Franssen et al., 2020; Kochs et al., 2023; Pimpini et al., 2022). Under the present incidental-viewing task, however, palatability did not emerge as a stable positive stimulus-linked representational dimension comparable to the shared calorie/health axis observed in occipitotemporal cortex.

### Limitations and future directions

Several limitations should be considered. The final sample was modest (n = 25, after excluding one participant who could not complete scanning and seven with incompatible slice-acquisition and phase-encoding parameters) and consisted exclusively of healthy young women within a narrow BMI range. The female-only design, which we adopted a priori to reduce variability associated with reported sex differences in neural responses to food cues, limits generalizability across sexes and precludes characterising between-sex differences. The modest sample size also reduces the precision and stability of the estimated effect sizes. This concern applies to the RSA effects generally, and is especially relevant for the more complex model-comparison analyses, such as the commonality partitions, where unique components were small and often not reliable after correction. Accordingly, the reported effect magnitudes should be interpreted cautiously and require independent replication.

Relatedly, because the sample comprised normative-weight young women, it is unclear how far the present representational profiles generalise across the BMI spectrum. Whether neural responses to food images differ with BMI is itself unsettled: a preregistered meta-analysis found no overall difference between individuals with obesity and lean individuals (Morys, García-García, & Dagher, 2023), and our own work has not found reliable BMI effects on food-cue responses (Pimpini et al., 2022). We therefore do not assume that the present profiles would differ in higher-BMI populations, and frame the perceived calorie/health representational structure identified here as a finding in a normative-weight sample whose generalisation across BMI remains an open question for future work.

Additionally, ratings were collected post-scan and may therefore reflect memory-based or reflective judgments rather than online valuation processes occurring during stimulus presentation. Because our design cannot determine the direction of any resulting bias, offline ratings could either weaken correspondence with state- and context-dependent regions or fail to capture online evaluative structure altogether. We therefore treat rating timing as one of several non-exclusive contributors to the absence of reliable effects in OFC and insula, rather than as a directional account of those nulls. For the stable, stimulus-linked structure in ventral visual cortex, this concern is less acute.

The rapid event-related design, short stimulus presentation times, and the orthogonal task, may have preferentially captured early perceptual and ventral stream representations, while reducing sensitivity to slower, integrative value signals typically associated with higher-order regions such as OFC, insula, and prefrontal cortex. Value computation and cognitive control processes often unfold over longer timescales and may require sustained attention, explicit evaluation, or decision engagement to become reliably expressed at the neural level. As a result, the present design may have biased representational sensitivity toward stimulus-driven visual representations at the expense of temporally extended valuation dynamics.

Although we complemented the ROI analyses with parcel-wise RSA, which provides finer spatial resolution within the same regions, both analyses rest on a fixed group parcellation and on representational structure that is consistent across participants. They may therefore remain insensitive to coding schemes that do not respect group-defined parcel boundaries, that vary in location across individuals, or that are distributed or dynamic, particularly in association and prefrontal regions. Approaches such as subject-specific searchlight or hyperalignment-based RSA would be better suited to recovering such patterns.

Future work could address these limitations by manipulating metabolic state (e.g., hunger vs satiety), varying task demands to explicitly engage subjective judgment or decision-related processing, and employing time-resolved or connectivity-based representational analyses to capture temporally evolving food-value signals.

## Conclusion

The present study shows that food-property information is reflected in occipitotemporal representational geometry during incidental food viewing. Visual model structure dominated early visual cortex, whereas perceived calorie, perceived health, and objective calorie category showed smaller but reliable effects in LOTC and VOTC. However, these effects did not resolve into separable calorie- or health-specific representations. Instead, residualized RSA and commonality analyses indicated that perceived calorie, perceived health, and objective calorie category contributed largely through shared representational structure. Thus, calorie-related effects in occipitotemporal cortex are best interpreted as part of a broader food-property axis, linking energy-density cues, perceived health, category structure, and learned food knowledge, rather than as an isolated representation of caloric magnitude.

## Acknowledgements

We thank Sanne Schins (MRI research technician at Scannexus) and Reja Debevc (student assistant at Maastricht University) for their help during participants’ testing. This study was financed by a VIDI grant of the Dutch Research Council (NWO) (016.165.356) awarded to Prof. dr. Anne Roefs and by starter grant of the Dutch Ministry of Education, Culture and Science (OCW) (SG2024-5) awarded to dr. Leonardo Pimpini.

## Declaration of generative AI and AI-assisted technologies in the writing process

During the preparation of this work the author(s) used claude.ai in order to improve readability of the manuscript. After using this tool/service, the author(s) reviewed and edited the content as needed and take(s) full responsibility for the content of the published article.

## CRediT authorship contribution statement

**Giuseppe Marrazzo**: Conceptualization, Methodology, Data Curation, Validation, Formal Analysis, Visualization, Software, Writing – original draft, Writing – review & editing. **Leonardo Pimpini**: Project administration, Conceptualization, Methodology, Investigation, Funding acquisition, Writing – original draft, Writing – review & editing. **Sarah Kochs:** Conceptualization, Methodology, Investigation, Software, Writing – review & editing. **Federico De Martino**: Conceptualization, Methodology, writing – review & editing. **Giancarlo Valente**: Software, Validation, Writing – review & editing. **Anne Roefs**: Funding acquisition, Conceptualization, Methodology, Supervision, Writing – review & editing.

