## Supplementary Material for "Representational structure of perceived food attributes in human occipitotemporal cortex"

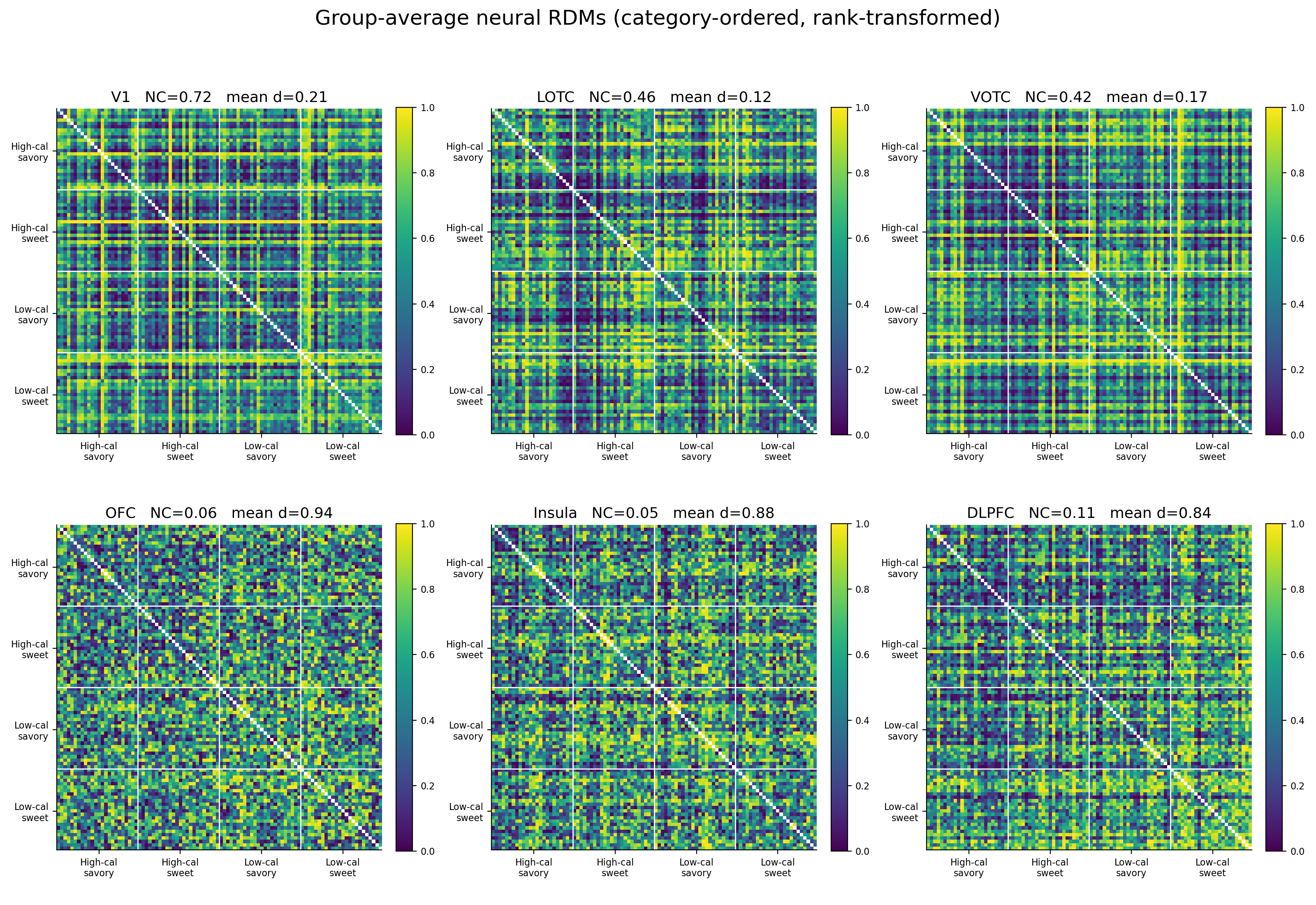

**Supplementary Figure S1. Group-average neural RDMs across regions of interest.**
Group-average neural representational dissimilarity matrices (RDMs) are shown for V1, lateral occipitotemporal cortex (LOTC), ventral occipitotemporal cortex (VOTC), orbitofrontal cortex (OFC), insula, and dorsolateral prefrontal cortex (DLPFC). For each participant and ROI, neural RDMs were computed from condition-level response patterns across the 96 food stimuli using correlation distance. RDMs were then averaged across participants for visualization and rank-transformed to emphasize relative dissimilarity structure. Stimuli are ordered by the factorial stimulus categories: high-calorie savory, high-calorie sweet, low-calorie savory, and low-calorie sweet. Warmer colors indicate greater neural dissimilarity between stimulus pairs. NC indicates the split-half neural RDM noise ceiling for each ROI, and mean d indicates the mean rank-transformed dissimilarity value. These group-average RDMs are shown for descriptive visualization only; all statistical inference was performed at the subject level. The visual and occipitotemporal ROIs showed more reliable stimulus-linked structure than OFC, insula, and DLPFC, where low noise ceilings indicate weaker stable neural RDM structure under the present task and analysis.

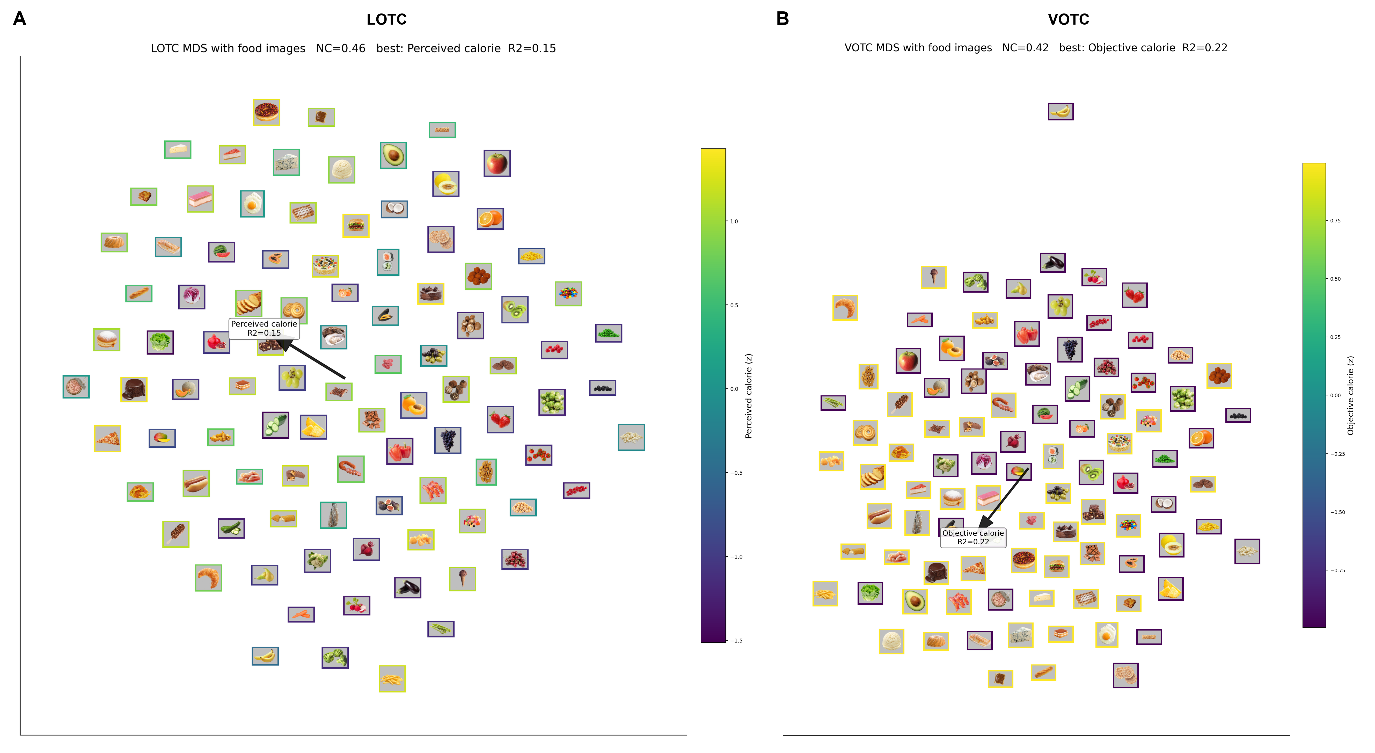

**Supplementary Figure S2. Neural MDS visualization of food-stimulus geometry in LOTC and VOTC.**
Two-dimensional non-metric multidimensional scaling (MDS) visualizations of the group-average neural representational dissimilarity matrices (RDMs) in lateral occipitotemporal cortex (LOTC; A) and ventral occipitotemporal cortex (VOTC; B). For each participant and ROI, condition-level response patterns were first estimated for each of the 96 food stimuli. Neural RDMs were then computed by calculating the correlation distance between all pairs of stimulus-specific response patterns within the ROI. For visualization, these subject-level neural RDMs were averaged across participants to obtain one group-average neural RDM per ROI. Non-metric MDS was applied to each group-average RDM to obtain a two-dimensional layout in which stimuli with more similar ROI-level response patterns are placed closer together and stimuli with more dissimilar response patterns are placed farther apart. Each thumbnail represents one food stimulus. Thumbnail positions were adjusted slightly after MDS to reduce visual overlap, while preserving the overall geometry of the embedding.

To aid interpretation of the MDS layouts, we fitted simple linear property gradients across the two-dimensional stimulus coordinates. For each candidate stimulus property, including perceived calorie, objective calorie category, palatability, savory/sweet category, and image-color properties, z-scored property values were regressed onto the two MDS coordinates. The property with the highest regression R² was selected for visualization in each ROI. Border colors indicate the value of this best-fitting property for each stimulus: perceived calorie in LOTC and objective calorie category in VOTC. Arrows indicate the direction of increasing property values in the MDS space. The reported R² values quantify how much variance in the corresponding stimulus property is captured by a linear gradient across the two-dimensional MDS layout. These R² values are descriptive visualization metrics and do not represent neural variance explained, RSA model–brain correlation strength, or statistical evidence for a model effect. All inferential analyses were performed at the subject level using RSA and permutation testing. NC indicates the split-half neural RDM noise ceiling for the corresponding ROI.

**Supplementary Table S1. Summary of significant whole-cortex parcel-wise RSA effects.**
The table reports bilateral Glasser parcels surviving FDR correction across the 179 analyzable bilateral parcels separately for each model or residualized control contrast (qFDR < .05). The bilateral H parcel did not yield valid group-level RSA estimates after grayordinate masking and was excluded from group statistics. Values in parentheses indicate mean Spearman rho across participants and the corresponding FDR-corrected q-value. Models and contrasts without FDR-surviving positive parcels are not shown. Group perceived-calorie, perceived-health, and perceived-palatability models were constructed from group-average z-scored ratings.

| **Model / contrast** | **N positive parcels** | **Positive parcels** |
| --- | --- | --- |
| Gabor | 17 | V2 (rho = 0.366, q < .001); V1 (rho = 0.355, q < .001); V3 (rho = 0.346, q < .001); V4 (rho = 0.296, q < .001); V8 (rho = 0.172, q < .001); PIT (rho = 0.101, q < .001); V3A (rho = 0.095, q < .001); V3CD (rho = 0.090, q < .001); VMV3 (rho = 0.064, q < .001); V3B (rho = 0.059, q = .003); LO1 (rho = 0.050, q = .015); IFJp (rho = 0.034, q = .021); MIP (rho = 0.034, q = .015); PEF (rho = 0.033, q = .007); IFJa (rho = 0.031, q = .039); POS2 (rho = 0.029, q = .003); PeEc (rho = 0.023, q = .015) |
| CORnet V4 | 34 | V2 (rho = 0.166, q < .001); V4 (rho = 0.145, q < .001); V1 (rho = 0.135, q < .001); V3 (rho = 0.135, q < .001); V8 (rho = 0.086, q < .001); V3A (rho = 0.062, q < .001); V3CD (rho = 0.055, q < .001); V3B (rho = 0.050, q < .001); LO1 (rho = 0.049, q < .001); LO2 (rho = 0.039, q < .001); V4t (rho = 0.038, q < .001); VMV3 (rho = 0.034, q < .001); PIT (rho = 0.031, q < .001); V7 (rho = 0.030, q < .001); VVC (rho = 0.028, q < .001); PH (rho = 0.028, q < .001); LO3 (rho = 0.024, q < .001); FFC (rho = 0.023, q < .001); IP0 (rho = 0.022, q < .001); PGp (rho = 0.020, q < .001); IPS1 (rho = 0.019, q < .001); VMV2 (rho = 0.018, q < .001); MT (rho = 0.014, q < .001); FST (rho = 0.014, q < .001); V6A (rho = 0.009, q = .003); PHA3 (rho = 0.009, q = .030); MST (rho = 0.008, q = .020); 8BM (rho = 0.008, q = .002); TPOJ3 (rho = 0.007, q = .040); MIP (rho = 0.007, q = .038); PreS (rho = 0.007, q = .007); PEF (rho = 0.007, q = .043); VIP (rho = 0.006, q = .038); LIPv (rho = 0.006, q = .022) |
| CORnet IT | 33 | V4 (rho = 0.130, q < .001); V2 (rho = 0.124, q < .001); V3 (rho = 0.111, q < .001); V1 (rho = 0.104, q < .001); V8 (rho = 0.085, q < .001); V3CD (rho = 0.056, q < .001); LO1 (rho = 0.052, q < .001); V3A (rho = 0.050, q < .001); V3B (rho = 0.048, q < .001); LO2 (rho = 0.046, q < .001); VMV3 (rho = 0.044, q < .001); VVC (rho = 0.041, q < .001); PIT (rho = 0.040, q < .001); PH (rho = 0.038, q < .001); V4t (rho = 0.038, q < .001); V7 (rho = 0.029, q < .001); LO3 (rho = 0.025, q < .001); IP0 (rho = 0.023, q < .001); FFC (rho = 0.023, q < .001); IPS1 (rho = 0.021, q < .001); FST (rho = 0.020, q < .001); PGp (rho = 0.020, q < .001); MT (rho = 0.018, q < .001); VMV2 (rho = 0.015, q < .001); PHA3 (rho = 0.012, q = .001); MST (rho = 0.009, q = .006); LIPv (rho = 0.009, q = .001); MIP (rho = 0.009, q = .029); VMV1 (rho = 0.008, q = .041); p32 (rho = 0.008, q = .020); V6A (rho = 0.007, q = .024); 8BM (rho = 0.006, q = .036); TPOJ3 (rho = 0.005, q = .021) |
| Group perceived calorie | 18 | VVC (rho = 0.034, q < .001); PH (rho = 0.027, q = .002); V8 (rho = 0.026, q < .001); FST (rho = 0.025, q = .001); V3CD (rho = 0.025, q < .001); LO1 (rho = 0.022, q < .001); VMV3 (rho = 0.021, q = .002); MST (rho = 0.021, q < .001); V4t (rho = 0.020, q = .013); LO3 (rho = 0.019, q = .007); MT (rho = 0.018, q = .031); PIT (rho = 0.017, q = .002); LO2 (rho = 0.014, q = .014); PHT (rho = 0.013, q = .018); TPOJ3 (rho = 0.012, q = .031); V4 (rho = 0.012, q = .004); PoI2 (rho = 0.012, q = .018); IP0 (rho = 0.012, q = .031) |
| Objective calorie | 30 | VVC (rho = 0.034, q < .001); PH (rho = 0.029, q < .001); V3CD (rho = 0.027, q < .001); V8 (rho = 0.025, q < .001); VMV3 (rho = 0.025, q < .001); LO1 (rho = 0.023, q < .001); LO3 (rho = 0.022, q < .001); V4t (rho = 0.018, q < .001); FST (rho = 0.018, q < .001); PIT (rho = 0.017, q < .001); FFC (rho = 0.016, q < .001); V4 (rho = 0.016, q < .001); IP0 (rho = 0.015, q < .001); LO2 (rho = 0.014, q < .001); MST (rho = 0.013, q < .001); MT (rho = 0.013, q = .006); PHT (rho = 0.012, q = .005); V3B (rho = 0.012, q < .001); V3A (rho = 0.012, q < .001); IPS1 (rho = 0.012, q < .001); PGp (rho = 0.012, q = .009); VMV2 (rho = 0.011, q = .002); V7 (rho = 0.010, q = .016); PHA3 (rho = 0.010, q = .001); PoI2 (rho = 0.009, q = .034); MIP (rho = 0.007, q = .038); V3 (rho = 0.006, q < .001); 47l (rho = 0.006, q = .045); TPOJ2 (rho = 0.006, q = .038); V2 (rho = 0.003, q = .018) |
| Group perceived health | 15 | V4t (rho = 0.038, q < .001); VVC (rho = 0.035, q < .001); PH (rho = 0.030, q < .001); MT (rho = 0.028, q = .002); FST (rho = 0.028, q < .001); V3CD (rho = 0.027, q < .001); LO1 (rho = 0.026, q < .001); LO3 (rho = 0.025, q = .001); LO2 (rho = 0.024, q < .001); MST (rho = 0.023, q < .001); VMV3 (rho = 0.021, q = .001); IP0 (rho = 0.018, q = .020); PIT (rho = 0.017, q = .005); V8 (rho = 0.015, q = .013); PHT (rho = 0.012, q = .015) |
| Savory/sweet category | 3 | LO2 (rho = 0.011, q = .043); PH (rho = 0.010, q = .020); FFC (rho = 0.007, q = .020) |
| Group perceived calorie controlling for Color | 15 | VVC (rho = 0.033, q < .001); V8 (rho = 0.028, q < .001); PH (rho = 0.027, q = .003); V3CD (rho = 0.025, q < .001); FST (rho = 0.024, q = .001); LO1 (rho = 0.022, q = .002); MST (rho = 0.020, q = .002); VMV3 (rho = 0.020, q = .003); V4t (rho = 0.020, q = .023); PIT (rho = 0.019, q < .001); LO3 (rho = 0.018, q = .032); V4 (rho = 0.016, q = .002); LO2 (rho = 0.014, q = .017); PHT (rho = 0.013, q = .023); PoI2 (rho = 0.012, q = .017) |
| Group perceived calorie controlling for CORnet V4 | 16 | VVC (rho = 0.033, q = .002); PH (rho = 0.026, q = .003); FST (rho = 0.025, q = .003); V3CD (rho = 0.021, q = .003); V8 (rho = 0.021, q = .002); MST (rho = 0.020, q = .002); VMV3 (rho = 0.019, q = .003); LO1 (rho = 0.019, q = .003); V4t (rho = 0.018, q = .031); LO3 (rho = 0.018, q = .015); MT (rho = 0.017, q = .048); PIT (rho = 0.015, q = .005); PHT (rho = 0.013, q = .022); LO2 (rho = 0.012, q = .047); PoI2 (rho = 0.012, q = .022); TPOJ3 (rho = 0.011, q = .048) |
| Group perceived calorie controlling for CORnet IT | 9 | VVC (rho = 0.028, q = .002); FST (rho = 0.022, q = .005); PH (rho = 0.021, q = .035); MST (rho = 0.020, q = .002); LO3 (rho = 0.015, q = .049); VMV3 (rho = 0.014, q = .035); LO1 (rho = 0.013, q = .041); PHT (rho = 0.013, q = .041); PoI2 (rho = 0.011, q = .041) |
| Group perceived calorie controlling for group perceived health | 10 | V2 (rho = 0.042, q = .001); V1 (rho = 0.032, q = .001); V3 (rho = 0.031, q = .001); IFSa (rho = 0.028, q = .026); V8 (rho = 0.026, q = .008); V4 (rho = 0.025, q = .004); PEF (rho = 0.022, q = .024); p9-46v (rho = 0.016, q = .037); 23c (rho = 0.015, q = .025); p32 (rho = 0.012, q = .046) |
| Group perceived health controlling for group perceived calorie | 2 | V4t (rho = 0.040, q = .029); MT (rho = 0.024, q = .041) |
| Objective calorie controlling for Color | 29 | VVC (rho = 0.033, q < .001); PH (rho = 0.029, q < .001); V3CD (rho = 0.027, q < .001); V8 (rho = 0.026, q < .001); VMV3 (rho = 0.024, q < .001); LO1 (rho = 0.023, q < .001); LO3 (rho = 0.021, q < .001); PIT (rho = 0.018, q < .001); V4t (rho = 0.018, q < .001); V4 (rho = 0.018, q < .001); FST (rho = 0.017, q < .001); FFC (rho = 0.016, q < .001); IP0 (rho = 0.015, q < .001); LO2 (rho = 0.014, q < .001); V3A (rho = 0.013, q < .001); MST (rho = 0.012, q < .001); PHT (rho = 0.012, q = .005); MT (rho = 0.012, q = .010); V3B (rho = 0.012, q < .001); IPS1 (rho = 0.012, q < .001); PGp (rho = 0.012, q = .013); V7 (rho = 0.010, q = .022); VMV2 (rho = 0.010, q = .005); PoI2 (rho = 0.009, q = .022); V3 (rho = 0.009, q < .001); PHA3 (rho = 0.009, q = .007); V2 (rho = 0.006, q < .001); VMV1 (rho = 0.006, q = .038); V1 (rho = 0.004, q = .006) |
| Objective calorie controlling for CORnet V4 | 27 | VVC (rho = 0.032, q < .001); PH (rho = 0.027, q < .001); V3CD (rho = 0.024, q < .001); VMV3 (rho = 0.023, q < .001); LO1 (rho = 0.021, q < .001); LO3 (rho = 0.021, q < .001); V8 (rho = 0.021, q < .001); FST (rho = 0.017, q < .001); V4t (rho = 0.016, q < .001); PIT (rho = 0.016, q < .001); FFC (rho = 0.014, q = .001); IP0 (rho = 0.014, q < .001); MST (rho = 0.012, q < .001); PHT (rho = 0.012, q = .006); MT (rho = 0.012, q = .010); LO2 (rho = 0.012, q < .001); IPS1 (rho = 0.011, q < .001); PGp (rho = 0.011, q = .020); VMV2 (rho = 0.010, q = .005); V3B (rho = 0.009, q = .007); PHA3 (rho = 0.009, q = .002); PoI2 (rho = 0.009, q = .036); V7 (rho = 0.009, q = .045); V3A (rho = 0.009, q = .007); V4 (rho = 0.008, q = .009); 47l (rho = 0.006, q = .048); TPOJ2 (rho = 0.006, q = .039) |
| Objective calorie controlling for CORnet IT | 19 | VVC (rho = 0.028, q < .001); PH (rho = 0.024, q < .001); V3CD (rho = 0.019, q < .001); VMV3 (rho = 0.019, q < .001); LO3 (rho = 0.019, q = .003); LO1 (rho = 0.016, q = .002); FST (rho = 0.015, q = .001); V8 (rho = 0.013, q = .008); V4t (rho = 0.013, q = .001); FFC (rho = 0.013, q = .003); IP0 (rho = 0.012, q = .002); PIT (rho = 0.012, q = .002); PHT (rho = 0.012, q = .006); MST (rho = 0.012, q < .001); MT (rho = 0.010, q = .047); IPS1 (rho = 0.009, q = .003); VMV2 (rho = 0.009, q = .011); PHA3 (rho = 0.008, q = .008); LO2 (rho = 0.007, q = .012) |
| Objective calorie controlling for group perceived health | 8 | V2 (rho = 0.038, q < .001); V1 (rho = 0.032, q < .001); V3 (rho = 0.026, q < .001); V3A (rho = 0.026, q = .001); V4 (rho = 0.023, q = .001); V8 (rho = 0.020, q < .001); VMV2 (rho = 0.015, q < .001); VMV1 (rho = 0.014, q = .046) |

**Supplementary Table 2. Grayordinate feature counts for predefined ROI parcel groups.**

The table lists the bilateral Glasser parcels included in the predefined ROI summaries shown in Figure 3A. Left and right HCP-MMP labels were collapsed into bilateral parcel IDs. Feature counts report the number of valid grayordinates retained for RSA after excluding non-finite, all-zero, or zero-variance features. Because the final analysis used all valid grayordinates, the number of valid grayordinates used is the effective feature count for each parcel. Values are shown as mean (minimum-maximum) across participants when they varied across subjects, or as a single value when constant.

| **ROI group** | **Parcel** | **Valid grayordinates used** |
| --- | --- | --- |
| V1 | V1 | 1618 |
| LOTC | LO1 | 80 |
| LOTC | LO2 | 83 |
| LOTC | LO3 | 125 |
| VOTC | FFC | 318 |
| VOTC | VVC | 250 |
| OFC | 11l | 323 |
| OFC | 13l | 199 |
| OFC | OFC | 680 |
| OFC | pOFC | 195.2 (187-196) |
| Insula | PoI2 | 333 |
| Insula | MI | 224 |
| Insula | AVI | 276 |
| Insula | AAIC | 139 |
| Insula | PoI1 | 317 |
| Insula | Ig | 189 |
| Insula | PI | 257 |
| DLPFC | 8Av | 406 |
| DLPFC | 8Ad | 481 |
| DLPFC | 8BL | 341 |
| DLPFC | 9p | 250 |
| DLPFC | 8C | 500 |
| DLPFC | p9-46v | 309 |
| DLPFC | 46 | 456 |
| DLPFC | a9-46v | 221 |
| DLPFC | 9-46d | 516 |
| DLPFC | 9a | 282 |
| DLPFC | i6-8 | 185 |
| DLPFC | s6-8 | 141 |

**Supplementary Table S3. Summary and rationale for RSA model RDMs and control analyses.**

The table summarizes the representational dissimilarity matrices (RDMs), derived behavioral variants, residualized RSA controls, and commonality analyses used to evaluate visual, subjective, categorical, and controlled accounts of neural representational structure. Continuous behavioral RDMs were constructed from within-participant z-scored ratings. Residualized RSA controls test partial associations in rank-transformed RDM space and are not interpreted as formal variance decompositions; formal unique and shared variance was assessed with the commonality analyses listed at the bottom of the table.

| **Model / analysis** | **Type** | **What it captures** | **How it was constructed** | **Interpretive role** |
| --- | --- | --- | --- | --- |
| **Gabor** | Low-level visual RDM | Orientation, spatial-frequency, and texture-like image structure. | Pairwise dissimilarities between Gabor feature representations of the 96 food images. | Positive effects indicate neural sensitivity to early image-computable visual structure. |
| **Color histogram** | Low-level visual RDM | Global colour-distribution similarity across images. | Pairwise dissimilarities between Lab colour-histogram feature vectors. | Used as a low-level visual control for colour-based stimulus similarity. |
| **CORnet V4** | High-level visual RDM | Intermediate visual object features approximating mid-level ventral-stream representations. | Pairwise dissimilarities between CORnet V4 feature representations. | Tests whether neural geometry follows mid-level visual structure. |
| **CORnet IT** | High-level visual RDM | Higher-level object/semantic visual structure approximating later ventral-stream representations. | Pairwise dissimilarities between CORnet IT feature representations. | Tests whether neural geometry follows high-level visual object structure. |
| **PercPal** | Subjective behavioral RDM | Group-level rating differences for palatability | Within-participant z-scored ratings were averaged across participants to obtain one consensus value per stimulus; pairwise stimulus differences were then computed. | Tests whether neural similarity reflects subjective tastiness/palatability structure. |
| **PercCal** | Subjective behavioral RDM | Group-level rating differences for perceived calorie content. | Within-participant z-scored ratings were averaged across participants to obtain one consensus value per stimulus; pairwise stimulus differences were then computed. | Primary subjective nutritional model; tests whether neural geometry reflects perceived calorie structure. |
| **PercHealth** | Subjective behavioral RDM | Group-level rating differences for perceived health value. | Within-participant z-scored ratings were averaged across participants to obtain one consensus value per stimulus; pairwise stimulus differences were then computed. | Tests whether neural geometry reflects perceived health structure. |
| **ObjCal** | Categorical RDM | Predefined high-calorie versus low-calorie category membership. | Binary category-mismatch RDM: pairs from different calorie categories were coded as more dissimilar than pairs from the same category. | Tests whether neural geometry follows objective calorie category rather than graded subjective calorie ratings. |
| **SavorySweet** | Categorical RDM | Predefined savory versus sweet category membership. | Binary category-mismatch RDM based on savory/sweet stimulus labels. | Tests whether neural geometry reflects the balanced savory/sweet category distinction. |
| **PercCal/Health/ObjCal \| visual controls** | Residualized RSA control | Test the neural association after removing visual-model structure. | Neural and model RDM vectors were rank-transformed and residualized with respect to a visual control RDM such as CORnet IT, CORnet V4, or Color; residual vectors were then correlated. | Positive effects indicate target model (ie PercCal) neural structure not explained by the specified visual model alone. |
| **PercCal \| PercHealth** | Residualized RSA control | Tests the neural association of perceived-calorie structure after removing the component linearly related to perceived-health structure. | Neural and PercCal RDM vectors were rank-transformed and residualized with respect to PercHealth residual vectors were then correlated. | Positive effects indicate that perceived-calorie structure remained associated with neural representational geometry after removing the component shared with perceived-health structure. |

Abbreviations: PercCal = perceived calorie; PercHealth = perceived health; PercPal = perceived palatability; ObjCal = objective calorie category.

**Supplementary Table S4. Bayesian evidence for ROI-level RSA effects in higher-order ROIs.**
Bayes factors were computed on Fisher-z-transformed ROI-level RSA correlations. Mean ρ and SEM are reported on the original correlation scale. *p*-values are uncorrected two-sided one-sample *t*-tests on the Fisher-z-transformed ROI-level effects. BF10 quantifies evidence for a non-zero effect relative to the null; BF01 quantifies evidence for the null relative to a non-zero effect. Evidence labels combine Bayes-factor strength with the sign of the mean Fisher-z effect. Group perceived-calorie, perceived-health, and perceived-palatability models were constructed from group-average z-scored ratings. Subject-own models were constructed separately for each participant from that participant’s own ratings.

| **ROI** | **Model / contrast** | **Mean ρ ± SEM** | ***t*(24)** | ***p*(two-sided)** | **BF10** | **BF01** | **Evidence** |
| --- | --- | --- | --- | --- | --- | --- | --- |
| OFC | Group perceived calorie | −0.0001 ± 0.0029 | −0.03 | .977 | 0.21 | 4.74 | Moderate null |
| OFC | Group perceived health | −0.0018 ± 0.0030 | −0.60 | .555 | 0.25 | 4.03 | Moderate null |
| OFC | Group perceived palatability | −0.0053 ± 0.0037 | −1.41 | .171 | 0.51 | 1.97 | Inconclusive |
| OFC | Group perceived calorie \| group perceived health | +0.0030 ± 0.0042 | +0.70 | .492 | 0.26 | 3.80 | Moderate null |
| OFC | Own perceived calorie | +0.0023 ± 0.0031 | +0.72 | .477 | 0.27 | 3.74 | Moderate null |
| OFC | Own perceived health | −0.0003 ± 0.0030 | −0.09 | .932 | 0.21 | 4.73 | Moderate null |
| OFC | Own perceived palatability | −0.0027 ± 0.0048 | −0.58 | .570 | 0.25 | 4.08 | Moderate null |
| Insula | Group perceived calorie | +0.0051 ± 0.0037 | +1.38 | .179 | 0.49 | 2.03 | Inconclusive |
| Insula | Group perceived health | +0.0055 ± 0.0038 | +1.46 | .157 | 0.54 | 1.85 | Inconclusive |
| Insula | Group perceived palatability | +0.0105 ± 0.0060 | +1.75 | .093 | 0.79 | 1.26 | Inconclusive |
| Insula | Group perceived calorie \| group perceived health | +0.0006 ± 0.0070 | +0.09 | .933 | 0.21 | 4.73 | Moderate null |
| Insula | Own perceived calorie | +0.0019 ± 0.0048 | +0.40 | .691 | 0.23 | 4.41 | Moderate null |
| Insula | Own perceived health | +0.0056 ± 0.0045 | +1.24 | .228 | 0.42 | 2.40 | Inconclusive |
| Insula | Own perceived palatability | +0.0063 ± 0.0057 | +1.10 | .283 | 0.36 | 2.76 | Inconclusive |
| DLPFC | Group perceived calorie | +0.0008 ± 0.0035 | +0.22 | .828 | 0.22 | 4.64 | Moderate null |
| DLPFC | Group perceived health | −0.0089 ± 0.0031 | −2.86 | .009 | 5.42 | 0.18 | Moderate negative |
| DLPFC | Group perceived palatability | −0.0136 ± 0.0070 | −1.94 | .064 | 1.05 | 0.95 | Inconclusive |
| DLPFC | Group perceived calorie \| group perceived health | +0.0172 ± 0.0067 | +2.57 | .017 | 3.09 | 0.32 | Moderate positive |
| DLPFC | Own perceived calorie | +0.0044 ± 0.0048 | +0.92 | .368 | 0.31 | 3.25 | Moderate null |
| DLPFC | Own perceived health | −0.0072 ± 0.0034 | −2.10 | .046 | 1.37 | 0.73 | Inconclusive |
| DLPFC | Own perceived palatability | −0.0114 ± 0.0075 | −1.52 | .142 | 0.58 | 1.72 | Inconclusive |

**Note.** Positive and negative evidence labels indicate the direction of the observed mean Fisher-z effect. BF10 and BF01 are reciprocal Bayes factors. Conventional descriptive thresholds were used: BF > 3 = moderate evidence; BF > 10 = strong evidence. The contrast “group perceived calorie | group perceived health” tests whether group perceived-calorie structure remains associated with neural representational geometry after removing the component linearly related to group perceived-health structure. Models are shown only for OFC, insula, and DLPFC, the higher-order ROIs examined in this Bayesian sensitivity analysis.

**Supplementary Table S5. ROI RSA using participant-specific behavioral RDMs.**
For each participant, subject-specific perceived-calorie, perceived-health, and perceived-palatability RDMs were constructed from that participant’s own ratings by z-scoring ratings across stimuli and computing pairwise absolute differences. Each subject-specific behavioral RDM was correlated with the same participant’s neural RDM within each pooled ROI. Values indicate mean Spearman ρ across participants ± SEM. Directional permutation p-values were computed on Fisher-z-transformed subject-level correlations. *q*within ROI indicates FDR correction across the three subject-specific behavioral models within each ROI. *q*by model indicates FDR correction across ROIs for the same model.

| **ROI** | **Subject-specific model** | **Mean ρ** | **SEM** | ***p*perm** | ***q*within ROI** | ***q*by model** |
| --- | --- | --- | --- | --- | --- | --- |
| V1 | Own perceived calorie | −0.004 | 0.004 | .841 | .999 | .841 |
| V1 | Own perceived health | −0.015 | 0.004 | .999 | .999 | .999 |
| V1 | Own perceived palatability | −0.021 | 0.010 | .980 | .999 | .985 |
| LOTC | Own perceived calorie | 0.029 | 0.007 | < .001 | < .001 | < .001 |
| LOTC | Own perceived health | 0.032 | 0.005 | < .001 | < .001 | < .001 |
| LOTC | Own perceived palatability | −0.018 | 0.009 | .966 | .966 | .985 |
| VOTC | Own perceived calorie | 0.030 | 0.006 | < .001 | < .001 | < .001 |
| VOTC | Own perceived health | 0.034 | 0.006 | < .001 | < .001 | < .001 |
| VOTC | Own perceived palatability | −0.022 | 0.010 | .985 | .985 | .985 |
| OFC | Own perceived calorie | 0.002 | 0.003 | .241 | .720 | .362 |
| OFC | Own perceived health | 0.000 | 0.003 | .541 | .720 | .811 |
| OFC | Own perceived palatability | −0.003 | 0.005 | .720 | .720 | .985 |
| Insula | Own perceived calorie | 0.002 | 0.005 | .344 | .344 | .413 |
| Insula | Own perceived health | 0.006 | 0.005 | .114 | .219 | .227 |
| Insula | Own perceived palatability | 0.006 | 0.006 | .146 | .219 | .875 |
| DLPFC | Own perceived calorie | 0.004 | 0.005 | .184 | .553 | .362 |
| DLPFC | Own perceived health | −0.007 | 0.003 | .978 | .978 | .999 |
| DLPFC | Own perceived palatability | −0.011 | 0.007 | .927 | .978 | .985 |

**Supplementary Table S6. Split-half consistency of group-level behavioral rating models.**
Split-half consistency was computed across rating participants for perceived calorie, perceived health, and perceived palatability. For each random split, ratings were z-scored within participant and averaged within each half of participants. Consistency was computed both for the stimulus-wise group rating vectors and for the corresponding absolute-difference RDMs used in RSA. Values indicate Fisher-z-averaged Spearman correlations across 10,000 random balanced splits; brackets indicate the 2.5th–97.5th percentile range across splits. Spearman-Brown-corrected values estimate the reliability of the full-sample group model.

| **Rating dimension** | **Vector split-half ρ** | **Vector SB-corrected ρ** | **RDM split-half ρ** | **RDM SB-corrected ρ** |
| --- | --- | --- | --- | --- |
| Perceived calorie | .949 [.933, .962] | .974 [.965, .981] | .938 [.921, .953] | .968 [.959, .976] |
| Perceived health | .948 [.930, .962] | .974 [.964, .981] | .942 [.924, .955] | .970 [.960, .977] |
| Perceived palatability | .773 [.675, .840] | .872 [.806, .913] | .532 [.387, .638] | .694 [.558, .779] |

Note. SB = Spearman-Brown corrected. Vector consistency reflects the reliability of the stimulus-wise group-average rating axis. RDM consistency reflects the reliability of the behavioral model RDM after converting ratings into pairwise absolute differences and is therefore the more directly relevant reliability estimate for RSA.

**Supplementary Table S7. ROI-level RSA using group-level model RDMs.**

Group-level RSA effects were computed separately within each pooled ROI. Values indicate mean Spearman ρ across participants ± SEM (n = 25). Directional permutation p-values were computed on Fisher-z-transformed subject-level correlations (50,000 sign-flip permutations). q within ROI indicates Benjamini-Hochberg FDR correction across all tested models and contrasts within each ROI. q by model indicates FDR correction across ROIs for the same model or contrast. q by model x ROI indicates global Benjamini-Hochberg FDR correction across all ROI × model/contrast tests. The vertical bar denotes a residualized RSA control (target model after removing the component linearly related to the specified control model).

| **ROI** | **Model / contrast** | **Mean ρ ± SEM** | **p perm** | **q within ROI** | **q by model** | **q by model x ROI** |
| --- | --- | --- | --- | --- | --- | --- |
| V1 | Gabor | +0.355 ± 0.012 | < .001 | < .001 | < .001 | < .001 |
| V1 | Color | −0.053 ± 0.012 | 1.000 | 1.000 | 1.000 | 1.000 |
| V1 | CORnet V4 | +0.135 ± 0.008 | < .001 | < .001 | < .001 | < .001 |
| V1 | CORnet IT | +0.104 ± 0.006 | < .001 | < .001 | < .001 | < .001 |
| V1 | Group perceived calorie | −0.007 ± 0.003 | .993 | 1.000 | .993 | 1.000 |
| V1 | Group perceived health | −0.026 ± 0.003 | 1.000 | 1.000 | 1.000 | 1.000 |
| V1 | Group perceived palatability | −0.024 ± 0.009 | .993 | 1.000 | .993 | 1.000 |
| V1 | Objective calorie | +0.003 ± 0.001 | .013 | .043 | .026 | .040 |
| V1 | Savory/sweet | −0.003 ± 0.002 | .952 | 1.000 | .952 | 1.000 |
| V1 | Group perceived calorie │ Gabor | −0.017 ± 0.003 | 1.000 | 1.000 | 1.000 | 1.000 |
| V1 | Group perceived calorie │ Color | −0.004 ± 0.003 | .934 | 1.000 | .934 | 1.000 |
| V1 | Group perceived calorie │ CORnet V4 | −0.015 ± 0.003 | 1.000 | 1.000 | 1.000 | 1.000 |
| V1 | Group perceived calorie │ CORnet IT | −0.026 ± 0.003 | 1.000 | 1.000 | 1.000 | 1.000 |
| V1 | Group perceived calorie │ group perceived health | +0.032 ± 0.005 | < .001 | < .001 | < .001 | < .001 |
| V1 | Group perceived calorie │ objective calorie | −0.016 ± 0.004 | .999 | 1.000 | .999 | 1.000 |
| V1 | Objective calorie │ Gabor | −0.003 ± 0.001 | .983 | 1.000 | .983 | 1.000 |
| V1 | Objective calorie │ Color | +0.004 ± 0.001 | < .001 | .003 | .002 | .003 |
| V1 | Objective calorie │ CORnet V4 | −0.004 ± 0.001 | .999 | 1.000 | .999 | 1.000 |
| V1 | Objective calorie │ CORnet IT | −0.012 ± 0.001 | 1.000 | 1.000 | 1.000 | 1.000 |
| V1 | Objective calorie │ group perceived health | +0.032 ± 0.003 | < .001 | < .001 | < .001 | < .001 |
| V1 | Objective calorie │ group perceived calorie | +0.014 ± 0.004 | < .001 | .003 | .002 | .002 |
| V1 | Group perceived health │ Gabor | −0.030 ± 0.003 | 1.000 | 1.000 | 1.000 | 1.000 |
| V1 | Group perceived health │ Color | −0.025 ± 0.003 | 1.000 | 1.000 | 1.000 | 1.000 |
| V1 | Group perceived health │ CORnet V4 | −0.036 ± 0.003 | 1.000 | 1.000 | 1.000 | 1.000 |
| V1 | Group perceived health │ CORnet IT | −0.047 ± 0.003 | 1.000 | 1.000 | 1.000 | 1.000 |
| V1 | Group perceived health │ group perceived calorie | −0.041 ± 0.005 | 1.000 | 1.000 | 1.000 | 1.000 |
| V1 | Group perceived health │ objective calorie | −0.041 ± 0.005 | 1.000 | 1.000 | 1.000 | 1.000 |
| LOTC | Gabor | +0.044 ± 0.017 | .008 | .010 | .016 | .025 |
| LOTC | Color | −0.004 ± 0.015 | .601 | .649 | .934 | .861 |
| LOTC | CORnet V4 | +0.051 ± 0.004 | < .001 | < .001 | < .001 | < .001 |
| LOTC | CORnet IT | +0.058 ± 0.006 | < .001 | < .001 | < .001 | < .001 |
| LOTC | Group perceived calorie | +0.028 ± 0.005 | < .001 | < .001 | < .001 | < .001 |
| LOTC | Group perceived health | +0.037 ± 0.006 | < .001 | < .001 | < .001 | < .001 |
| LOTC | Group perceived palatability | −0.022 ± 0.009 | .988 | .988 | .993 | 1.000 |
| LOTC | Objective calorie | +0.030 ± 0.004 | < .001 | < .001 | < .001 | < .001 |
| LOTC | Savory/sweet | +0.006 ± 0.002 | .006 | .008 | .019 | .022 |
| LOTC | Group perceived calorie │ Gabor | +0.028 ± 0.005 | < .001 | < .001 | < .001 | < .001 |
| LOTC | Group perceived calorie │ Color | +0.029 ± 0.005 | < .001 | < .001 | < .001 | < .001 |
| LOTC | Group perceived calorie │ CORnet V4 | +0.026 ± 0.004 | < .001 | < .001 | < .001 | < .001 |
| LOTC | Group perceived calorie │ CORnet IT | +0.019 ± 0.004 | < .001 | < .001 | < .001 | < .001 |
| LOTC | Group perceived calorie │ group perceived health | −0.007 ± 0.008 | .800 | .831 | .800 | 1.000 |
| LOTC | Group perceived calorie │ objective calorie | +0.007 ± 0.005 | .087 | .103 | .497 | .194 |
| LOTC | Objective calorie │ Gabor | +0.030 ± 0.004 | < .001 | < .001 | < .001 | < .001 |
| LOTC | Objective calorie │ Color | +0.030 ± 0.004 | < .001 | < .001 | < .001 | < .001 |
| LOTC | Objective calorie │ CORnet V4 | +0.028 ± 0.004 | < .001 | < .001 | < .001 | < .001 |
| LOTC | Objective calorie │ CORnet IT | +0.022 ± 0.004 | < .001 | < .001 | < .001 | < .001 |
| LOTC | Objective calorie │ group perceived health | +0.006 ± 0.005 | .130 | .146 | .195 | .284 |
| LOTC | Objective calorie │ group perceived calorie | +0.012 ± 0.004 | .005 | .008 | .010 | .018 |
| LOTC | Group perceived health │ Gabor | +0.036 ± 0.006 | < .001 | < .001 | < .001 | < .001 |
| LOTC | Group perceived health │ Color | +0.037 ± 0.006 | < .001 | < .001 | < .001 | < .001 |
| LOTC | Group perceived health │ CORnet V4 | +0.033 ± 0.006 | < .001 | < .001 | < .001 | < .001 |
| LOTC | Group perceived health │ CORnet IT | +0.026 ± 0.006 | < .001 | < .001 | < .001 | < .001 |
| LOTC | Group perceived health │ group perceived calorie | +0.024 ± 0.009 | .006 | .008 | .037 | .022 |
| LOTC | Group perceived health │ objective calorie | +0.024 ± 0.009 | .006 | .008 | .037 | .022 |
| VOTC | Gabor | +0.044 ± 0.014 | .001 | .002 | .004 | .005 |
| VOTC | Color | +0.018 ± 0.018 | .164 | .185 | .934 | .346 |
| VOTC | CORnet V4 | +0.030 ± 0.003 | < .001 | < .001 | < .001 | < .001 |
| VOTC | CORnet IT | +0.039 ± 0.004 | < .001 | < .001 | < .001 | < .001 |
| VOTC | Group perceived calorie | +0.028 ± 0.005 | < .001 | < .001 | < .001 | < .001 |
| VOTC | Group perceived health | +0.030 ± 0.006 | < .001 | < .001 | < .001 | < .001 |
| VOTC | Group perceived palatability | −0.018 ± 0.012 | .917 | .917 | .993 | 1.000 |
| VOTC | Objective calorie | +0.032 ± 0.004 | < .001 | < .001 | < .001 | < .001 |
| VOTC | Savory/sweet | +0.007 ± 0.002 | < .001 | < .001 | .001 | < .001 |
| VOTC | Group perceived calorie │ Gabor | +0.026 ± 0.005 | < .001 | < .001 | < .001 | < .001 |
| VOTC | Group perceived calorie │ Color | +0.027 ± 0.005 | < .001 | < .001 | < .001 | < .001 |
| VOTC | Group perceived calorie │ CORnet V4 | +0.026 ± 0.005 | < .001 | < .001 | < .001 | < .001 |
| VOTC | Group perceived calorie │ CORnet IT | +0.021 ± 0.004 | < .001 | < .001 | < .001 | < .001 |
| VOTC | Group perceived calorie │ group perceived health | +0.002 ± 0.007 | .372 | .386 | .555 | .701 |
| VOTC | Group perceived calorie │ objective calorie | +0.004 ± 0.005 | .224 | .242 | .497 | .466 |
| VOTC | Objective calorie │ Gabor | +0.031 ± 0.004 | < .001 | < .001 | < .001 | < .001 |
| VOTC | Objective calorie │ Color | +0.031 ± 0.004 | < .001 | < .001 | < .001 | < .001 |
| VOTC | Objective calorie │ CORnet V4 | +0.030 ± 0.004 | < .001 | < .001 | < .001 | < .001 |
| VOTC | Objective calorie │ CORnet IT | +0.026 ± 0.004 | < .001 | < .001 | < .001 | < .001 |
| VOTC | Objective calorie │ group perceived health | +0.014 ± 0.005 | .007 | .009 | .020 | .022 |
| VOTC | Objective calorie │ group perceived calorie | +0.016 ± 0.004 | < .001 | < .001 | .002 | .002 |
| VOTC | Group perceived health │ Gabor | +0.030 ± 0.006 | < .001 | < .001 | < .001 | < .001 |
| VOTC | Group perceived health │ Color | +0.030 ± 0.006 | < .001 | < .001 | < .001 | < .001 |
| VOTC | Group perceived health │ CORnet V4 | +0.028 ± 0.006 | < .001 | < .001 | < .001 | < .001 |
| VOTC | Group perceived health │ CORnet IT | +0.024 ± 0.006 | < .001 | < .001 | < .001 | < .001 |
| VOTC | Group perceived health │ group perceived calorie | +0.013 ± 0.008 | .063 | .074 | .189 | .164 |
| VOTC | Group perceived health │ objective calorie | +0.013 ± 0.008 | .063 | .074 | .189 | .164 |
| OFC | Gabor | +0.004 ± 0.006 | .262 | .819 | .314 | .517 |
| OFC | Color | −0.005 ± 0.006 | .758 | .819 | .934 | .999 |
| OFC | CORnet V4 | 0.000 ± 0.002 | .567 | .819 | .681 | .828 |
| OFC | CORnet IT | 0.000 ± 0.002 | .557 | .819 | .668 | .820 |
| OFC | Group perceived calorie | 0.000 ± 0.003 | .513 | .819 | .616 | .770 |
| OFC | Group perceived health | −0.002 ± 0.003 | .726 | .819 | 1.000 | .986 |
| OFC | Group perceived palatability | −0.005 ± 0.004 | .914 | .914 | .993 | 1.000 |
| OFC | Objective calorie | 0.000 ± 0.003 | .451 | .819 | .542 | .731 |
| OFC | Savory/sweet | +0.001 ± 0.002 | .409 | .819 | .541 | .724 |
| OFC | Group perceived calorie │ Gabor | 0.000 ± 0.003 | .527 | .819 | .632 | .783 |
| OFC | Group perceived calorie │ Color | 0.000 ± 0.003 | .481 | .819 | .578 | .743 |
| OFC | Group perceived calorie │ CORnet V4 | 0.000 ± 0.003 | .510 | .819 | .612 | .770 |
| OFC | Group perceived calorie │ CORnet IT | 0.000 ± 0.003 | .504 | .819 | .605 | .770 |
| OFC | Group perceived calorie │ group perceived health | +0.003 ± 0.004 | .242 | .819 | .485 | .497 |
| OFC | Group perceived calorie │ objective calorie | −0.001 ± 0.004 | .574 | .819 | .688 | .830 |
| OFC | Objective calorie │ Gabor | 0.000 ± 0.003 | .458 | .819 | .687 | .732 |
| OFC | Objective calorie │ Color | +0.001 ± 0.003 | .437 | .819 | .524 | .731 |
| OFC | Objective calorie │ CORnet V4 | 0.000 ± 0.003 | .448 | .819 | .672 | .731 |
| OFC | Objective calorie │ CORnet IT | 0.000 ± 0.003 | .444 | .819 | .666 | .731 |
| OFC | Objective calorie │ group perceived health | +0.002 ± 0.004 | .258 | .819 | .310 | .516 |
| OFC | Objective calorie │ group perceived calorie | +0.001 ± 0.004 | .418 | .819 | .501 | .724 |
| OFC | Group perceived health │ Gabor | −0.002 ± 0.003 | .727 | .819 | 1.000 | .986 |
| OFC | Group perceived health │ Color | −0.002 ± 0.003 | .707 | .819 | 1.000 | .986 |
| OFC | Group perceived health │ CORnet V4 | −0.002 ± 0.003 | .722 | .819 | 1.000 | .986 |
| OFC | Group perceived health │ CORnet IT | −0.002 ± 0.003 | .714 | .819 | 1.000 | .986 |
| OFC | Group perceived health │ group perceived calorie | −0.003 ± 0.004 | .789 | .819 | 1.000 | 1.000 |
| OFC | Group perceived health │ objective calorie | −0.003 ± 0.004 | .789 | .819 | 1.000 | 1.000 |
| Insula | Gabor | −0.002 ± 0.007 | .625 | .703 | .625 | .889 |
| Insula | Color | −0.006 ± 0.007 | .778 | .841 | .934 | 1.000 |
| Insula | CORnet V4 | −0.003 ± 0.002 | .862 | .862 | .862 | 1.000 |
| Insula | CORnet IT | −0.002 ± 0.002 | .840 | .862 | .840 | 1.000 |
| Insula | Group perceived calorie | +0.005 ± 0.004 | .087 | .148 | .175 | .194 |
| Insula | Group perceived health | +0.006 ± 0.004 | .078 | .148 | .157 | .187 |
| Insula | Group perceived palatability | +0.010 ± 0.006 | .047 | .148 | .282 | .141 |
| Insula | Objective calorie | +0.005 ± 0.003 | .053 | .148 | .079 | .144 |
| Insula | Savory/sweet | 0.000 ± 0.003 | .451 | .543 | .541 | .731 |
| Insula | Group perceived calorie │ Gabor | +0.005 ± 0.004 | .085 | .148 | .170 | .194 |
| Insula | Group perceived calorie │ Color | +0.005 ± 0.004 | .080 | .148 | .159 | .187 |
| Insula | Group perceived calorie │ CORnet V4 | +0.005 ± 0.004 | .082 | .148 | .164 | .190 |
| Insula | Group perceived calorie │ CORnet IT | +0.006 ± 0.004 | .079 | .148 | .157 | .187 |
| Insula | Group perceived calorie │ group perceived health | +0.001 ± 0.007 | .462 | .543 | .555 | .732 |
| Insula | Group perceived calorie │ objective calorie | +0.001 ± 0.005 | .395 | .508 | .593 | .720 |
| Insula | Objective calorie │ Gabor | +0.005 ± 0.003 | .054 | .148 | .108 | .146 |
| Insula | Objective calorie │ Color | +0.005 ± 0.003 | .051 | .148 | .076 | .144 |
| Insula | Objective calorie │ CORnet V4 | +0.005 ± 0.003 | .051 | .148 | .103 | .144 |
| Insula | Objective calorie │ CORnet IT | +0.006 ± 0.003 | .052 | .148 | .103 | .144 |
| Insula | Objective calorie │ group perceived health | +0.002 ± 0.004 | .315 | .501 | .315 | .615 |
| Insula | Objective calorie │ group perceived calorie | +0.002 ± 0.005 | .342 | .508 | .501 | .660 |
| Insula | Group perceived health │ Gabor | +0.006 ± 0.004 | .078 | .148 | .157 | .187 |
| Insula | Group perceived health │ Color | +0.006 ± 0.004 | .074 | .148 | .148 | .184 |
| Insula | Group perceived health │ CORnet V4 | +0.006 ± 0.004 | .073 | .148 | .146 | .184 |
| Insula | Group perceived health │ CORnet IT | +0.006 ± 0.004 | .066 | .148 | .133 | .170 |
| Insula | Group perceived health │ group perceived calorie | +0.002 ± 0.007 | .384 | .508 | .767 | .706 |
| Insula | Group perceived health │ objective calorie | +0.002 ± 0.007 | .384 | .508 | .767 | .706 |
| DLPFC | Gabor | +0.018 ± 0.008 | .017 | .229 | .025 | .052 |
| DLPFC | Color | +0.005 ± 0.014 | .371 | .975 | .934 | .701 |
| DLPFC | CORnet V4 | +0.001 ± 0.002 | .408 | .975 | .612 | .724 |
| DLPFC | CORnet IT | +0.003 ± 0.002 | .145 | .830 | .218 | .314 |
| DLPFC | Group perceived calorie | +0.001 ± 0.004 | .417 | .975 | .616 | .724 |
| DLPFC | Group perceived health | −0.009 ± 0.003 | .996 | .997 | 1.000 | 1.000 |
| DLPFC | Group perceived palatability | −0.014 ± 0.007 | .967 | .997 | .993 | 1.000 |
| DLPFC | Objective calorie | −0.001 ± 0.002 | .736 | .997 | .736 | .986 |
| DLPFC | Savory/sweet | +0.003 ± 0.002 | .154 | .830 | .308 | .328 |
| DLPFC | Group perceived calorie │ Gabor | 0.000 ± 0.004 | .470 | .975 | .632 | .732 |
| DLPFC | Group perceived calorie │ Color | +0.001 ± 0.004 | .447 | .975 | .578 | .731 |
| DLPFC | Group perceived calorie │ CORnet V4 | +0.001 ± 0.004 | .420 | .975 | .612 | .724 |
| DLPFC | Group perceived calorie │ CORnet IT | 0.000 ± 0.004 | .467 | .975 | .605 | .732 |
| DLPFC | Group perceived calorie │ group perceived health | +0.017 ± 0.007 | .008 | .220 | .024 | .026 |
| DLPFC | Group perceived calorie │ objective calorie | +0.003 ± 0.005 | .249 | .975 | .497 | .504 |
| DLPFC | Objective calorie │ Gabor | −0.002 ± 0.002 | .776 | .997 | .931 | 1.000 |
| DLPFC | Objective calorie │ Color | −0.002 ± 0.002 | .743 | .997 | .743 | .986 |
| DLPFC | Objective calorie │ CORnet V4 | −0.001 ± 0.002 | .741 | .997 | .889 | .986 |
| DLPFC | Objective calorie │ CORnet IT | −0.002 ± 0.002 | .792 | .997 | .951 | 1.000 |
| DLPFC | Objective calorie │ group perceived health | +0.007 ± 0.004 | .051 | .461 | .102 | .144 |
| DLPFC | Objective calorie │ group perceived calorie | −0.003 ± 0.004 | .819 | .997 | .819 | 1.000 |
| DLPFC | Group perceived health │ Gabor | −0.009 ± 0.003 | .996 | .997 | 1.000 | 1.000 |
| DLPFC | Group perceived health │ Color | −0.009 ± 0.003 | .996 | .997 | 1.000 | 1.000 |
| DLPFC | Group perceived health │ CORnet V4 | −0.009 ± 0.003 | .996 | .997 | 1.000 | 1.000 |
| DLPFC | Group perceived health │ CORnet IT | −0.010 ± 0.003 | .997 | .997 | 1.000 | 1.000 |
| DLPFC | Group perceived health │ group perceived calorie | −0.019 ± 0.006 | .997 | .997 | 1.000 | 1.000 |
| DLPFC | Group perceived health │ objective calorie | −0.019 ± 0.006 | .997 | .997 | 1.000 | 1.000 |
